# Bacterial community dynamics explain carbon mineralization and assimilation in soils of different land-use history

**DOI:** 10.1101/2022.02.16.480692

**Authors:** Samuel E. Barnett, Nicholas D. Youngblut, Daniel H. Buckley

**Author notes:** Corresponding author: Daniel H. Buckley.

## Abstract

Soil dwelling microorganisms are key players in the terrestrial carbon cycle, driving both the degradation and stabilization of soil organic matter. Bacterial community structure and function vary with respect to land-use, yet the ecological drivers of this variation remain poorly described and difficult to predict. We conducted a multi-substrate DNA-stable isotope probing experiment across cropland, old-field, and forest habitats to link carbon mineralization dynamics with the dynamics of bacterial growth and carbon assimilation. We tracked the movement of ^13^C derived from five distinct carbon sources as it was assimilated into bacterial DNA over time. We show that carbon mineralization, community composition, and carbon assimilation dynamics all differed with respect to land-use. We also show that microbial community dynamics affect carbon assimilation dynamics and are predictable from soil DNA content. Soil DNA yield is easy to measure and it predicts microbial community dynamics linked to soil carbon cycling.

**Originality-Significance Statement:** Soil dwelling microorganisms are key players in the terrestrial carbon cycle, driving both the degradation and stabilization of soil organic matter. Microbial communities vary with respect to land-use, but we still have an incomplete understanding of how variation in community structure links to variation in community function. DNA stable isotope probing (DNA-SIP) is a high-resolution method that can identify specific microbial taxa that assimilate carbon *in situ*. We conducted a large-scale multi-substrate DNA-SIP experiment to explore differences in bacterial activity across land-use regimes. We show that microbial community dynamics vary with land-use, that these dynamics are linked to soil carbon cycling, and that they are predicted from easily measured soil properties.

## Introduction

Soils contain a considerable portion of the terrestrial carbon (C) pool (Jobbágy and Jackson, 2000), and land-use affects soil capacity to process and store C. For example, conversion of natural land (*e.g.* forest and grassland) to cropland results in loss of soil organic matter (SOM) and reduced rates of C sequestration (Deng *et al*., 2016), while restoration (*e.g.* reforestation) increases soil C stocks (Post and Kwon, 2000). Soil C loss (*e.g.* mineralization to CO_2_) and stabilization (*e.g.* incorporation into SOM) result from the activities of soil dwelling microorganisms (Schimel and Schaeffer, 2012), with microbial biomass itself being a major component of SOM (Kallenbach *et al*., 2016; Liang *et al*., 2019). Recognizing the key role of microorganisms, recent soil C-cycle models, such as MIMICS (Wieder *et al*., 2014), CORPSE (Sulman *et al*., 2014), and MEND (Wang *et al*., 2015), include microbial processes in predicting soil C dynamics. However, C-cycle models do not account for changes in bacterial community structure and function due to land-use, because the mechanisms by which land-use alters microbial contributions to soil C-cycling remain uncertain.

Land-use changes alter the diversity of soil microbial communities (Lauber *et al*., 2008; Drenovsky *et al*., 2010; Jangid *et al*., 2011; Szoboszlay *et al*., 2017; Barnett *et al*., 2020) and these effects can persist for decades (Buckley and Schmidt, 2001, 2003; Jangid *et al*., 2011). Changes in community structure driven by land-use can also alter soil processes. For example, Tardy *et al*. found a positive correlation between soil respiration and bacterial richness across land-use (Tardy *et al*., 2015). Degens *et al*. demonstrated that soil catabolic diversity varies with management intensity (Degens *et al*., 2000). In addition, Malik *et al*. found that bacterial C-use efficiency correlates with land-use intensification (Malik *et al*., 2018). In general, management intensity exhibits a negative correlation with SOM, microbial biomass, and the size of the bacterial community (Degens *et al*., 2000; Nsabimana *et al*., 2004; Fornasier *et al*., 2014; Zhang *et al*., 2016; Albright *et al*., 2020). However, we still lack a firm grasp on how habitat and land-use characteristics regulate the activities of individual microbes within complex communities. Such knowledge would be useful in determining how microbial dynamics contribute to system level processes.

A major source of uncertainty in predicting microbial contributions to soil C-cycling is that traits realized *in situ* do not often correspond to trait potential predicted from metagenomic sequences or from analyses performed with cultured isolates. For example, a large proportion of the soil bacterial community is not metabolically active at any given time (Lennon and Jones, 2011; Blagodatskaya and Kuzyakov, 2013). Hence, the presence or absence of a given gene or even a given microbe might not infer activity. In addition, the traits that regulate microbial growth and survival in soil are far different from those that govern growth and survival in aqueous batch culture (Li *et al*., 2019), and the phenotypes exhibited in aqueous culture can differ dramatically from those exhibited in the soil matrix (Jansson and Hofmockel, 2018). As a result, our ability to predict microbial contributions to soil C-cycling are complicated by disjunction between the potential activities of microbes, as predicted from genome sequences or from phenotypes observed in culture, and the realized activities that microbes manifest *in situ*.

We conducted two concurrent experiments to examine how land-use influences realized bacterial activity in the C-cycle (Fig. 1). Experiments were performed with soils from a series of cropland, old-field, and restored forest sites, representing the three most common land-use types in the Finger Lakes region of NY, USA (Table S1; Fig. S1). We tracked C mineralization, bacterial C assimilation, and bacterial community dynamics by using experiments that employed five distinct C sources: xylose (a monomer of hemicellulose), an amino acid mixture (monomers of proteins), vanillin (an aromatic derivative of lignin), cellulose (a structural polymer in plant cell walls), and palmitic acid (a fatty acid constituent of plant cell membranes). We chose these substrates because they are readily derived from decomposing plant litter, their metabolism requires diverse pathways, and they range widely in bioavailability as determined by solubility and hydrophobicity (Barnett *et al*., 2021). In the first experiment (‘SIP Microcosms’, Fig. S2), performed at one set of sites, we performed multi-substrate stable isotope probing (SIP) to track C through the microbial food web. In the second experiment (‘Enrichment Microcosms’, Fig. S3), performed with a network of 10 locations (three sites each location) in a 50 km region, we examined bacterial community dynamics in response to the input of each C source. We hypothesized that bacterial contributions to soil C-cycling are not defined strictly by catabolic potential as encoded in the genome, but that they are instead determined habitat characteristics that influence growth dynamics. We predicted that discrete bacterial taxa found in two or more land-use types would exhibit land-use specific patterns of growth and carbon assimilation. We further hypothesized that realized participation in the C-cycle corresponds with bacterial growth dynamics, being driven by interactions between resource availability and bacterial life history traits (*i.e.,* bacterial allocation of energy to growth, resource acquisition, and survival).

**Figure 1:**
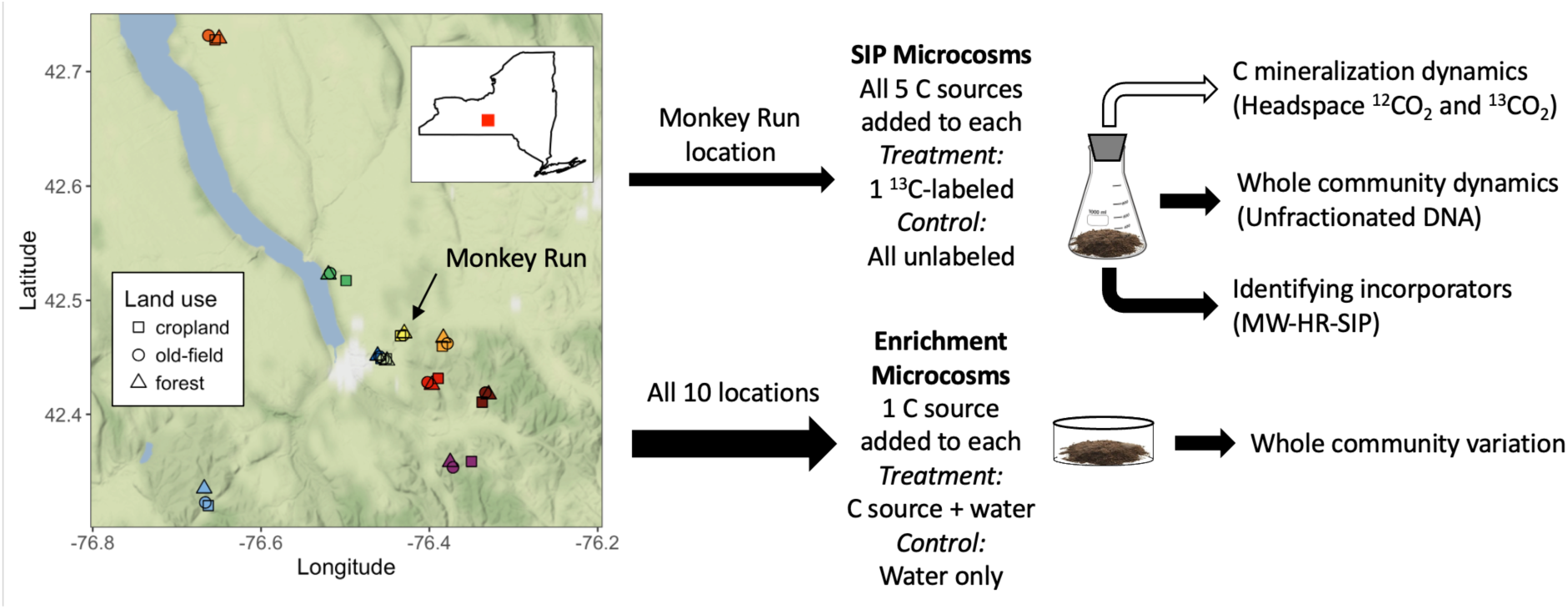
Overview of the experimental design which includes two separate soil microcosm experiments. SIP Microcosms were made with soil from cropland, old-field, and forest sites at the Monkey Run location and supplied with a mixture of all five C sources. SIP Microcosms were used to measure C mineralization dynamics, bacterial community dynamics, and to identify bacterial OTUs that assimilated ^13^C from each C source. Enrichment Microcosms were made using soils from each land-use from each of the 10 locations. Enrichment Microcosms were used to observe bacterial community change in response to each C source addition. These parallel microcosm experiments were designed to complement each other in resolution and scale. More detailed diagrams of these experiments are provided in Figs. S2 and S3.

## Results

### Carbon mineralization dynamics

We examined variation in C-cycling with respect to land-use in SIP microcosms by measuring C mineralization dynamics for the five C sources described above (see also Methods and Supplemental Methods). Variation in C mineralization rates across time were compared using linear mixed effects modeling. Cumulative C mineralized was compared across land-use using ANOVA. C mineralization dynamics varied significantly with respect to land-use, with cropland soils being most different from old-field and forest soils (Figs. 2, S4 – S6; Table S2 – S3). Total C mineralization rates (^12^CO_2_ + ^13^CO_2_) differed significantly with respect to land-use (Table S2; Fig. S4), with cropland soils reaching maximal mineralization rates 1 to 2 days later than old-field and forest soils (Fig. S4). Mineralization rates in cropland soils also fell more rapidly over time than other land-use types (Fig. S4). As a result, cropland soils mineralized significantly less C than the other land-use types (Table S3; Fig. S5). In addition, old-field soils had low mineralization rates during days 2 – 3 (Fig. S4), and as a result these soils mineralized significantly less C than forest soils (Fig. S5). These results reveal that the three land-use types had different C-mineralization dynamics and these differences were tied to significant variation in cumulative C mineralized from soils.

**Figure 2:**
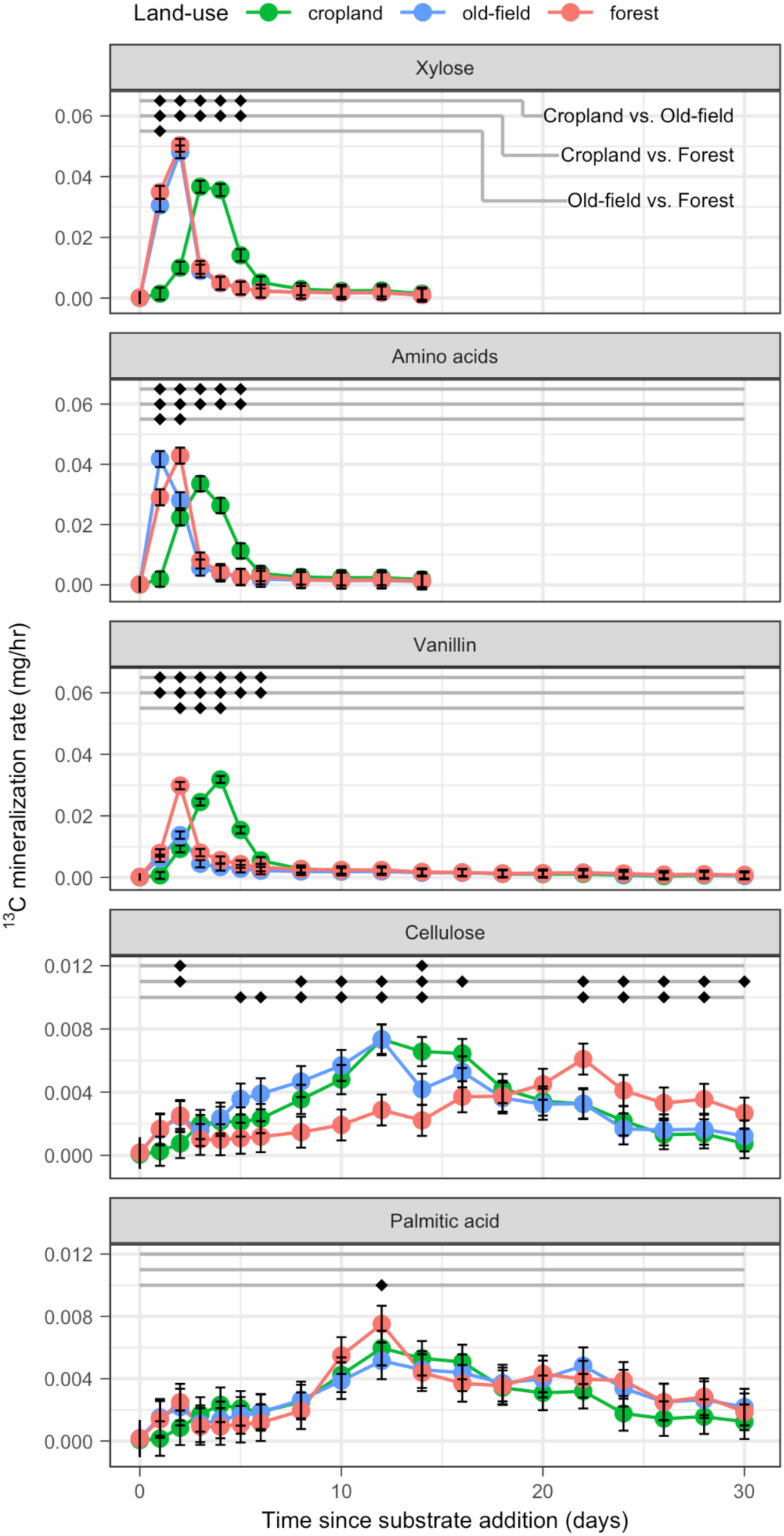
Carbon mineralization dynamics vary across land-use. Diamonds at the top of each panel indicate significant pairwise differences between land-use types at discrete points in time (adjusted *p-*value < 0.05). Each horizontal line indicates a different land-use comparison as identified in the top row. Error bars indicate standard error.

C mineralization rates for the five individual C sources (^13^CO_2_ only) varied with respect to time and its interaction with land-use (Fig. 2; Table S2). The dynamics of xylose mineralization were comparable in old-field and forest soils, though xylose mineralization in cropland soil exhibited a significant lag relative to the other soils (Fig. 2). Despite the difference in mineralization dynamics for xylose, land-use type did not affect cumulative xylose mineralization (Fig. S6). Amino acids and vanillin exhibited mineralization dynamics similar to xylose (Fig. 2), except that cropland soils mineralized significantly more vanillin-C than old-field or forest soils, and more amino acid-C than old-field soils (Fig. S6). In addition, the maximal rate of vanillin mineralization was significantly lower in old-field soils relative to forest soils and this reduction caused old-field soils to mineralize significantly less vanillin-C than forest soils (Figs. 2, S6). For cellulose, we saw similar mineralization dynamics in cropland and old-filed sites, as mineralization of cellulose-C in these soils reached a maximum on day 12 and both soils mineralized similar amounts of cellulose-C. In contrast, forest soils exhibited significantly lower cellulose mineralization rates on days 8 - 14 (Fig. 2), and significantly reduced cellulose-C mineralization relative to old-field soils (Fig. S6). Finally, mineralization dynamics and cumulative mineralization of palmitic acid did not differ significantly between the three soils in most respects (Figs. 2, S6). The results show that C mineralization dynamics, mineralization rates, and total C mineralized differ with respect to C source, land-use type, and the interaction of these two factors.

### Bacterial community dynamics in response to carbon addition

We assessed bacterial community dynamics in SIP Microcosm soils and Enrichment Microcosm soils after C addition. We evaluated community dynamics as a function of the DNA yield for initial field samples used to establish soil microcosms. In SIP Microcosms we assessed bacterial community dynamics using ‘unfractionated DNA’ (*i.e.,* DNA extracted from soils and sequenced directly without isopycnic centrifugation, see Fig. S2). We hypothesized that initial community size (*i.e,* bacterial biomass per g soil) would influence community dynamics in response to C addition, and that variation in soil DNA content might explain variation in community dynamics with respect to land use type. Bacterial biomass varies significantly in relation to land-use intensity (Nsabimana *et al*., 2004; Fornasier *et al*., 2014; Zhang *et al*., 2016; Albright *et al*., 2020). We used DNA yield as a proxy for microbial biomass, since soil DNA yield correlates strongly with soil microbial biomass and with cell numbers (Blagodatskaya *et al*., 2003; Joergensen and Emmerling, 2006; Blagodatskaya and Kuzyakov, 2013; Fornasier *et al*., 2014; Spohn *et al*., 2016).

In the SIP Microcosms, bacterial community alpha diversity (OTU richness, Shannon’s index, and Pielou’s evenness) varied with respect to land-use, time, and their interaction (Table S4). All soils experienced a drop in diversity within three days of C addition, but the effect was most dramatic in cropland soil (Figs. S7, S8), which had the lowest initial DNA yield. Alpha diversity changed less over time in forest and old-field soils than in cropland soils, with forest soils tending to have lower diversity than old-field soils (Figs. S7, S8). A similar pattern was observed for beta diversity (Bray-Curtis dissimilarity, unweighted and weighted UniFrac distance; Figs. 3, S9). Land-use explained most variation in community composition, but both time, and the interaction of land-use and time, also explained significant variation (PERMANOVA; Table 1). While bacterial communities from all land-use regimes changed over time in response to C addition, cropland communities exhibited the greatest change (Figs. 3, S9).

**Figure 3:**
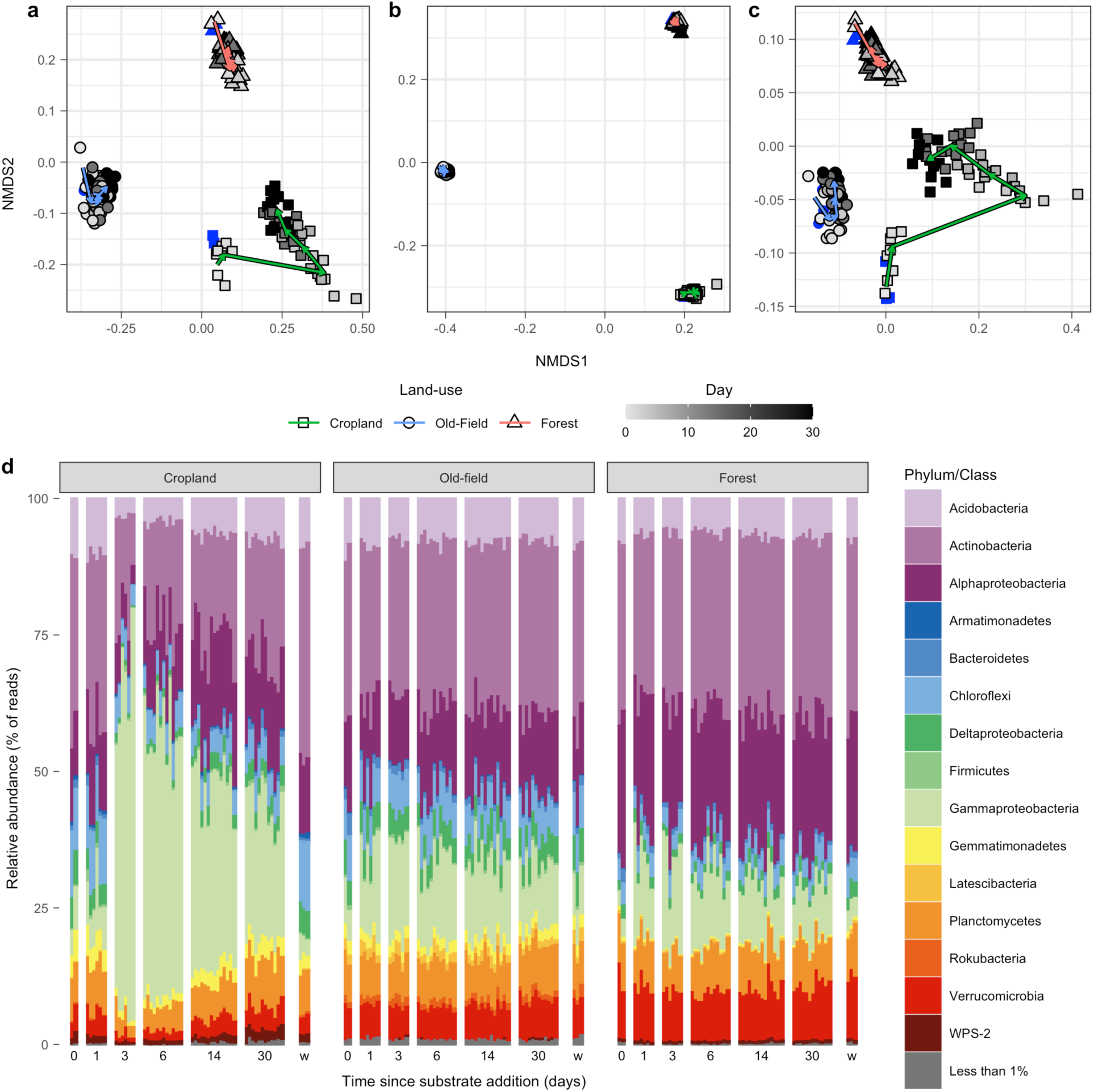
Bacterial communities in SIP Microcosms (unfractionated DNA) varied significantly across land-use, time since C addition (day), and their interaction (PERMANOVA; Table 1). These factors all explained significant variation in community composition using A) Bray-Curtis dissimilarity, B) unweighted UniFrac distance, and C) weighted UniFrac distance. Symbols indicate land-use, fill intensity indicates time, as defined in the legend, and arrows indicate shift in land-use centroids over time. Blue points represent the water only control microcosms sampled on day 30. D) Large changes in cropland soils over time can be attributed at least in part to dramatic expansion in the differential abundance of *Gammaproteobacteria* within 3 days of C addition. Note that the order *Betaproteobacteriales* is classified within the class *Gammaproteobacteria* (as of SILVA version 132). Each individual column represents a separate microcosm, grouped by sampling timepoint. The ‘w’ column indicates the water only control microcosms sampled on day 30.

**Table 1:**
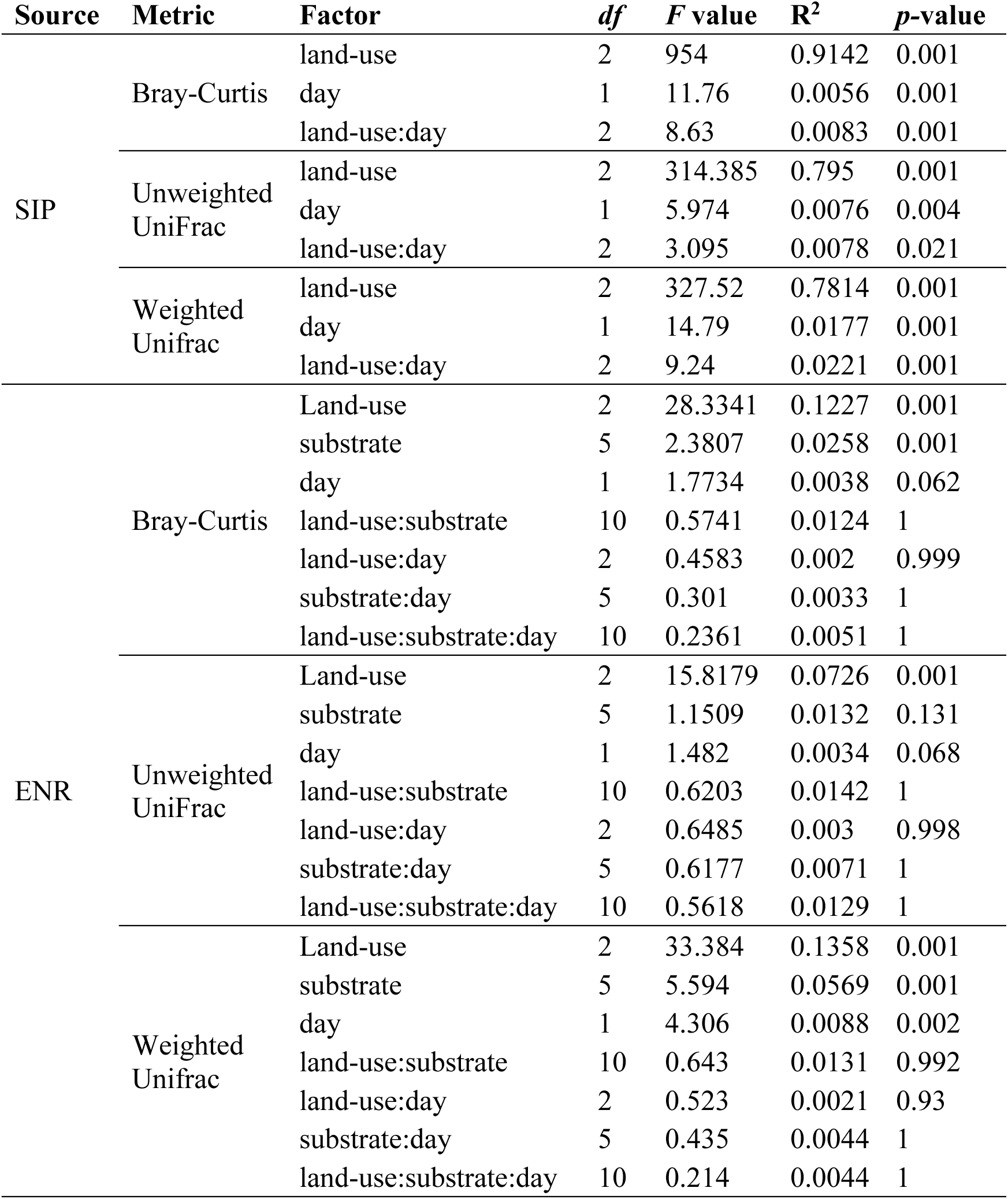
Land-use drives most variation in bacterial communities from SIP Microcosms (SIP) and Enrichment Microcosms (ENR), but communities also vary significantly with time (day; in both experiments) and substrate identity (in Enrichment Microcosms). Results are from PERMANOVA assessed using unfractionated DNA from soil microcosms.

Beta diversity within soils from the Enrichment Microcosms was also explained by land-use and time, but C source also had a significant effect (Table 1). Land-use explained most variation in bacterial beta diversity (Table 1). C source also explained significant variation in beta diversity as measured by Bray-Curtis dissimilarity and weighted UniFrac distance. Time explained significant variation in beta diversity only as measured by weighted UniFrac distance (PERMANOVA; Table 1). Variation in beta diversity within 2 days of adding amino acids or vanillin was correlated negatively with initial DNA yield (Pearson’s correlation; Fig. 4C; Table S5), indicating that the addition of these C sources caused a rapid change in community composition when DNA yield was low and less change in community composition when DNA yield was high. Similar correlations were observed for the other substrates at early time points, but these results were not significant.

**Figure 4:**
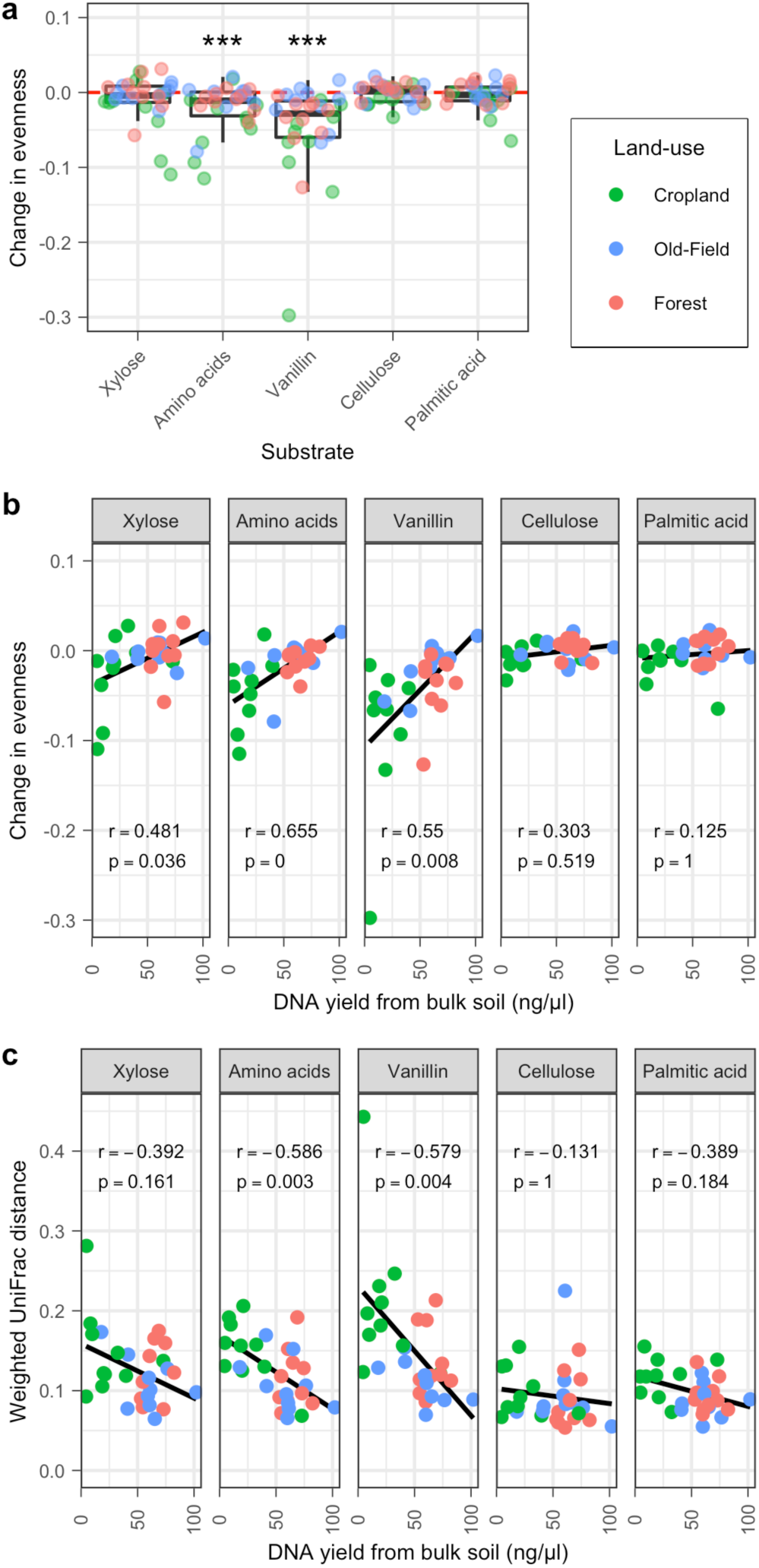
Change in community structure correlates with initial DNA yield from soil when soluble C sources are added to Enrichment Microcosms (*n* = 30 sites). A) Addition of amino acids or vanillin caused significant reduction in community evenness relative to water only controls (Pielou’s evenness; one-sample, one-sided Wilcoxon test, *** adjusted *p-*value < 0.001). B) Soils with small initial microbial populations exhibited greater reduction in evenness when soluble, but not insoluble, C sources were added. C) A similar relationship is observed for beta diversity (weighted UniFrac distance) for soils to which vanillin and amino acids were added. Lines calculated from linear regression, values indicate Pearson’s r and Benjamini-Hochberg adjusted *p-*value. Values are determined at early time points (*i.e.* xylose, amino acid, vanillin: day 2; cellulose and palmitic acid: day 14) between Enrichment Microcosm treatments that received C source addition and water only controls.

The evenness of bacterial communities in Enrichment Microcosms also declined significantly within two days of adding amino acids or vanillin (Wilcoxon test; Fig. 4A; Table S5). These changes in evenness correlated with initial DNA yield across all land use types (Pearson’s correlation; Fig. 4B; Table S5), indicating that evenness declined most when DNA yield was low. Xylose treatment exhibited a similar trend. However, no such correlation was observed for cellulose or palmitic acid treatments. With the exception of vanillin, these correlations were no longer significant at later time points, suggesting transient dynamics (Figs. S10; Table S5).

### Bacterial C assimilation varied by C source and land-use

We assessed differences in bacterial C assimilation across land-use using multiple-window high-resolution DNA-SIP (MW-HR-SIP) (Youngblut *et al*., 2018b, 2018a). This method allowed us to identify bacterial operational taxonomic units (OTU; clustered at 97% sequence identity) that assimilated ^13^C into their DNA. A total of 11331 OTUs passed sparsity filtering and 1452 of these exhibited significant ^13^C incorporation into their DNA (*i.e.* “incorporators”; supplemental data). There were 616 incorporators detected in cropland soil, 790 in old-field soil, and 574 in forest soil. Of these incorporators, 1055 were uniquely ^13^C-labeled in one land-use type, 266 were ^13^C-labeled in two land-use types, and 131 were ^13^C-labeled in all three land-use types (Fig. 5A). Of the 1452 incorporators observed, a total of 873 were present in all three land-use types (*i.e.,* passing sparsity filtering in at least one time point), but the fact that only 131 of these were ^13^C-labeled in all three land-use types indicates that patterns of ^13^C-assimilation were driven more by site identity than by OTU identity. This pattern, in which most OTUs are present in all three land-use types, but they are ^13^C-labeled in only one land-use type, was observed for all five C sources (Fig. S11).

**Figure 5:**
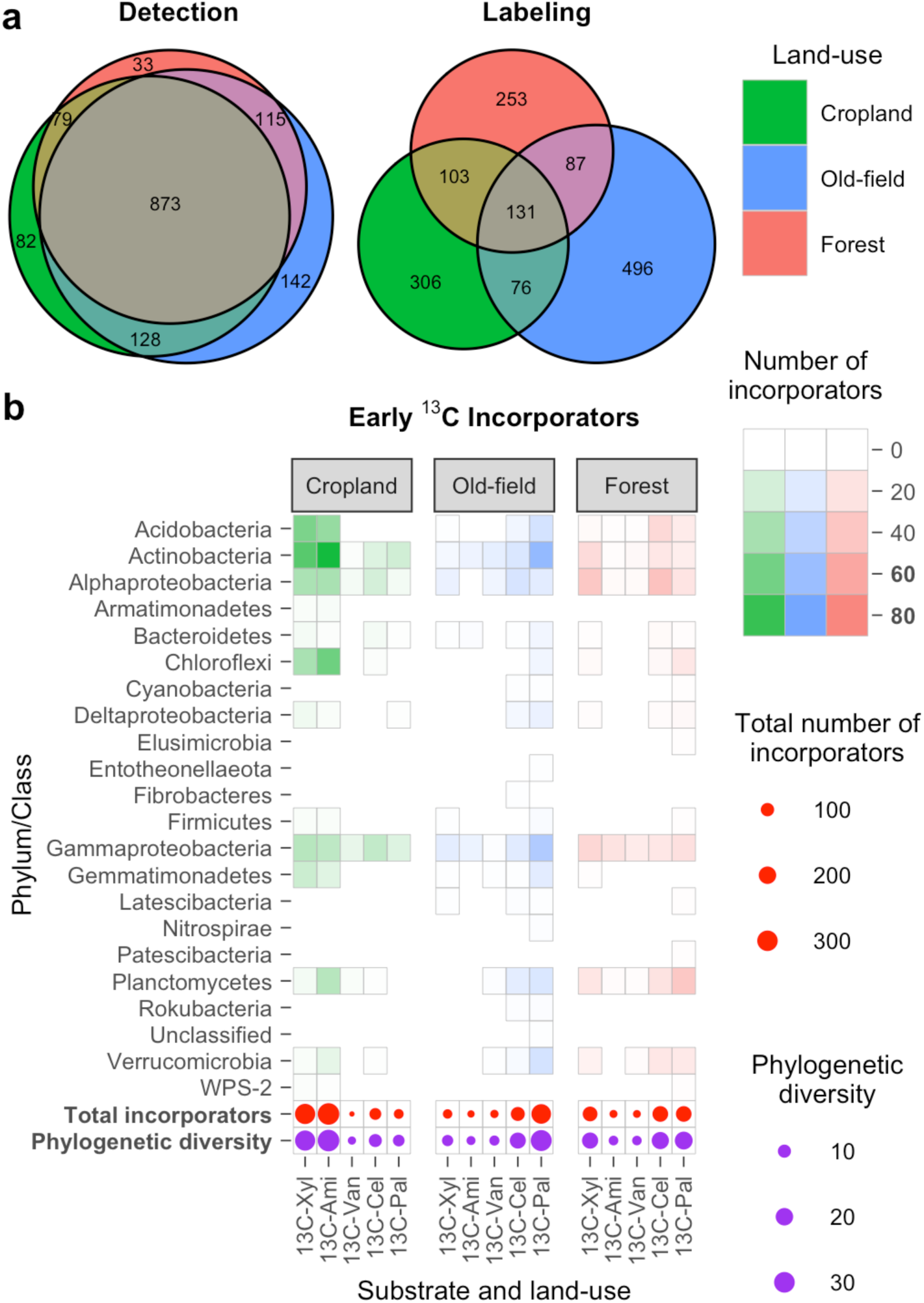
Patterns of C assimilation varied with respect to land-use. A) Among bacterial OTUs that assimilated ^13^C, most were observed in all land use types (Detection), but their ^13^C assimilation was driven by land-use (Labeling). B) During early time points (*i.e.* before peak mineralization rate for each substrate; xylose and amino acids: days 1 and 3; vanillin: day 6; cellulose and palmitic acid: days 6 and 14), incorporator diversity is higher for soluble C sources added to cropland soils, for insoluble C sources added to old-field and forest soils. Shading indicates the number of incorporators classified in each phylum/class. Total numbers and Faith’s phylogenetic diversity of the incorporators of each substrate indicated by circle sizes. Note that the order *Betaproteobacteriales* is classified within the class *Gammaproteobacteria* (as of SILVA version 132). Substrate names are abbreviated: 13C-Xyl = xylose, 13C-Ami = amino acids, 13C-Van = vanillin, 13C-Cel = cellulose, 13C-Pal = palmitic acid. Late incorporators shown in Fig. S10.

Incorporator diversity (richness, phylogenetic diversity, and taxonomic diversity) varied with respect to land-use and C source when assessed for early time points (Fig. 5B). Time points were defined as ‘early’ or ‘late’ based on the time of peak C mineralization for each C source. Early assimilation was defined as days 1 and 3 for xylose and amino acids, day 6 for vanillin, and days 6 and 14 for cellulose and palmitic acid. Early incorporators are those most likely to acquire C by direct assimilation from the C source. The diversity of early incorporators assimilating C from xylose and amino acids was highest in cropland soils (Fig. 5B), which had low DNA yields. Conversely, the diversity of early incorporators assimilating C from cellulose and palmitic acid was higher in old-field and forest soils than in cropland (Fig. 5B). The diversity of early incorporators assimilating C from vanillin was low in all three land-use types.

The pattern of assimilation was different for late incorporators (Fig. S12). Late time points were defined as those that occurred after the day peak C mineralization was observed for each C source (days 6 and 14 for xylose and amino acids, days 14 and 30 for vanillin, and day 30 for cellulose and palmitic acid). Late incorporators were likely to include taxa that acquire C by direct assimilation as well as those that assimilate C secondary to prior processing by other microbes. The diversity of late incorporators assimilating C from cellulose and palmitic acid was higher in forest soil than either old-field or cropland soils, and late incorporators of C from vanillin were higher in both forest and old-field soils than in cropland soil (Fig. S12). The diversity of late incorporators assimilating C from amino acids was highest in old-field soils (Fig. S12).

We used Enrichment Microcosms to examine OTUs that were enriched (as measured by differential abundance) by the addition of individual C sources. Enrichment Microcosms included soils from across 10 locations for each land-use. A total of 3638 OTUs from the Enrichment Microcosms passed sparsity filtering (representing at least 5 reads in at least 25% of the samples). Of these OTUs, 36 were enriched by C addition in at least one land-use type (cropland = 28, forest = 5, and old-field = 19; Supplemental Data), and 32 of these were also ^13^C incorporators in the SIP Microcosms (Supplemental Data). For 24 of these 32 OTUs, the C source that elicited a significant differential abundance (in Enrichment Microcosms) also elicited significant ^13^C-assimilation (in SIP Microcosms). These 32 OTUs belonged to four phyla (*Actinobacteria* = 8, *Bacteroidetes* =1, *Fimicutes* = 4, *Proteobacteria* = 23) and most were members of the family *Burkholderiaceae* (16 OTUs; Supplemental Data).

## Discussion

We show that variation in C-mineralization across land-use is coupled to bacterial growth dynamics and patterns of C assimilation. Patterns of C assimilation were particularly land-use specific as illustrated by the fact that 873 active taxa were common to all three land-use types but only 15% of these were consistently ^13^C-labeled across all land-use types (Fig. 5A). This simple yet notable finding adds to the growing body of evidence that bacterial occupancy, and thus metabolic potential, is not sufficient to predict bacterial activity in soil (Lennon and Jones, 2011; Aanderud *et al*., 2015; Fanin and Bertrand, 2016; Sorensen and Shade, 2020). One possible explanation for this result is that local dissemination may introduce microbes into non-optimal habitats where their activity is restricted by edaphic factors or competition from indigenous organisms (Tecon and Or, 2017). Such homogenizing dispersal has been identified as major driver of bacterial community assembly in the region that contains these sites (Barnett *et al*., 2020). In addition, dormancy and resuscitation, which themselves are likely to be associated with land-use specific edaphic features, may influence patterns of C assimilation (Lennon and Jones, 2011; Blagodatskaya and Kuzyakov, 2013).

We propose that microbial biomass quantity, as assessed here by DNA yield, may be an important ecological control on microbial processing of soil carbon. DNA yield, which correlates with microbial biomass (Blagodatskaya *et al*., 2003; Joergensen and Emmerling, 2006; Blagodatskaya and Kuzyakov, 2013; Fornasier *et al*., 2014; Spohn *et al*., 2016), is typically lower in cropland soil than in old-field and forest soil (Table S1). This result is unsurprising since intense agricultural management typically coincides with loss of microbial biomass (Nsabimana *et al*., 2004; Joergensen and Emmerling, 2006; Fornasier *et al*., 2014; Zhang *et al*., 2016; Albright *et al*., 2020). We hypothesize that DNA yield defines resource demand (*i.e.,* large population = high demand, small population = low demand) and interacts with resource supply to regulate C mineralization and assimilation dynamics in soil (Fig. 6). While this hypothesis requires further testing, we enumerate below how it can explain the results we observed.

**Figure 6:**
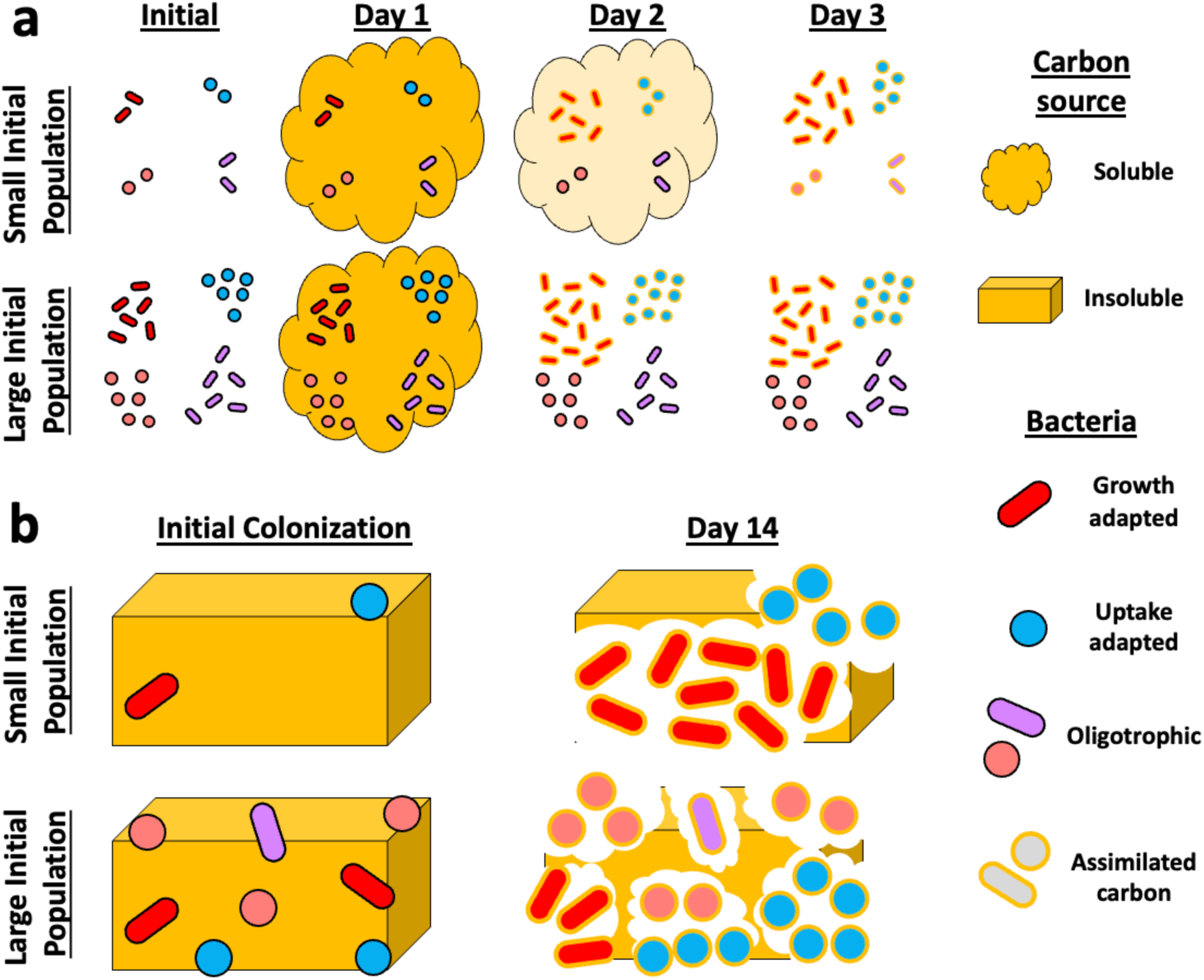
We hypothesize that initial microbial population size affects bacterial growth and assimilation dynamics. When the initial microbial population is small, new C inputs exceed initial demand leading to slow overall assimilation and mineralization of the C. As the populations expand fueled by the C and the C sources are depleted, demand eventually exceeds supply. When the initial microbial population is large, demand rapidly exceeds supply of new C inputs resulting in rapid assimilation and mineralization of the C. A) When soluble resources are added to water filled pore space high C source bioavailability generally favors microbes with the capability for highly dynamic or rapid growth. However, when initial population is small there is a window of time when supply exceeds demand and diverse microbes can assimilate C. A large initial population causes demand to exceed supply, imposing strict competition for the limited resources and short C residence time due to rapid assimilation. Such competition may cause C assimilation to be driven by high uptake rate as opposed to high growth rate. B) Insoluble resources do not diffuse through water filled pore space, and mineralization depends on colonization. A small initial population causes lower encounter rates, which provides an advantage to highly dynamic microbes that grow rapidly to monopolize a resource patch. A large initial population causes higher encounter rates, which blocks any one population from dominating the resource patch and favors microbes that compete best under conditions of resource limitation and microbe-microbe interactions (*e.g.,* auxotrophy).

First, in the SIP Microcosms, cropland soil exhibited a significant delay in the mineralization of highly bioavailable C (from xylose, amino acids, and vanillin), and this resulted in less overall C mineralization relative to old-field and forest soils (Figs. 2, S5). The mineralization rate of highly bioavailable C is a function of per cell activity and the number of cells present (Stenström *et al*., 1998; Blagodatskaya and Kuzyakov, 2013). Hence, soils with large initial microbial populations (*e.g.,* old-field and forest soils) can rapidly achieve maximal mineralization rates, while soils with small initial populations (*e.g.,* cropland soil) require a period of microbial growth before maximal mineralization rates are achieved (Fig. 6). The time required for growth would result in a lag before peak C mineralization is achieved. We expect growth lag to have less impact on the mineralization dynamics of low bioavailability C sources (*e.g.,* cellulose and palmitic acid) because mineralization of insoluble C is limited by the colonization of solid surfaces and by the limits of extracellular enzyme kinetics. Hence, we would expect the effect of microbial biomass quantity to have greater influence on the mineralization dynamics of highly bioavailable forms of C than on insoluble and surface associated C.

Many lines of evidence indicate that microbial biomass quantity influences bacterial growth dynamics and that these growth dynamics influence organic matter degradation. Highly bioavailable C is mineralized in the early stages of litter degradation (Berg *et al*., 1982; Berg, 2000; Don and Kalbitz, 2005), and delays in soil respiration during the early stages of litter degradation have been attributed to reduced microbial load (Albright *et al*., 2020). In addition, initial microbial load has been proposed to explain differences in decomposition rates across soil types (Rinkes *et al*., 2014). Finally, the mineralization of highly bioavailable C coincides with rapid changes in bacterial community composition and is linked to changes in bacterial C assimilation over time (Pepe-Ranney *et al*., 2016; Barnett *et al*., 2021),.

Second, cropland soil had more taxa assimilating C from high bioavailability sources at early time points than old-field and forest soils (Fig. 5B). This distinction can be explained by competition dynamics associated with highly bioavailable C sources (Kuzyakov and Blagodatskaya, 2015). When the demand for a resource exceeds the supply (*e.g.,* old-field and forest soils), high competition pressure will channel resources towards taxa that have adaptations maximizing resource acquisition. Such high competition for soluble organic matter has been observed in forest soils where microbial biomass is high (Goldfarb *et al*., 2011). However, when supply of a pulsed resource exceeds initial demand (*e.g.,* cropland soil), low competition will result in longer resource residence time, thereby allowing more taxa to access available resources (Fig. 6). Under these conditions, we would expect many microbes to grow rapidly while resources remain in excess, with growth terminating when resources are exhausted. Low competition coupled with high resource availability can thus explain the ‘boom-bust’ population dynamics apparent in the SIP Microcosms (Figs. 3, S13). Such ‘boom-bust’ growth can result in decreased community evenness and large changes in beta diversity as we see in the unfractionated DNA from SIP Microcosms, most prominently with cropland soil (Figs. S7-S9). Further, in the Enrichment Microcosms, we found correlations between beta diversity, evenness, and DNA yield (Fig. 4), suggesting that low DNA yield predicts communities that change dramatically when C is added to soils. Similar ‘boom-bust’ growth dynamics have been observed previously when highly bioavailable C sources are added to agricultural soils (Barnett *et al*., 2021). These observations suggest adaptive tradeoffs between those traits that favor rapid growth and the traits that favor efficient nutrient acquisition, as previously described (Fierer *et al*., 2007; Roller and Schmidt, 2015; Ramin and Allison, 2019; Malik *et al*., 2020; Barnett *et al*., 2021).

The most dynamic OTUs fell within the *Burkholderiacea* (genera of *Burkholderia-Caballeronia-Paraburkholderia*) and *Pseudomonas,* both of which include prominent ruderal strategists (Barnett *et al*., 2021), and are classified to the *Gammaproteobacteria* (as of SILVA version 132). These taxa assimilated ^13^C from high bioavailability C sources across all three land-use types (Supplemental Data), yet they increased far more dramatically in relative abundance in cropland soils than in old-field and forest soils (Figs. 3, S13). Population expansion of these taxa likely drove changes in both alpha and beta diversity observed in the SIP Microcosm soils following C addition (Table 1, S4; Figs. 3, S7, S8, S9).

Third, we found that early incorporators of ^13^C from insoluble substrates (cellulose and palmitic acid) were less diverse in cropland soils than in old-field and forest soils (Fig. 5B). Low bioavailability C sources like cellulose and palmitic acid are present in particulate form and their assimilation requires surface colonization and complex spatial interactions. Surface colonization in turn is determined by encounter rates and by expansion of populations across the particulate surface and through the particulate’s interior. Cell density dependent effects influence particulate organic matter degradation by bacteria in marine environments (Ebrahimi *et al*., 2019). We predict that encounter rates will be depressed when initial microbial populations are small, allowing initial colonists to have fewer competitors. Priority effects allow fast-growing initial colonizers to monopolize resources and prevent growth of secondary colonizers (Fukami, 2015; de Meester *et al*., 2016). When initial microbial populations are large, encounter rates will be enhanced allowing many bacterial cells to initially colonize particulate resources and leading to greater levels of competition (Fig. 6). It is also possible that low encounter rates favor the establishment of prototrophic organisms that have a full complement of biosynthetic pathways sufficient for growth, while high encounter rates favor auxotrophic organisms whose metabolic activities depend on products supplied by spatially co-localized organisms. In agricultural soils, microbes that have initial access to C from cellulose have been shown to be less auxotrophic than those microbes that access C at later time points (Wilhelm *et al*., 2021).

We found that relatively few OTUs in the Enrichment Microcosms responded consistently in differential abundance (*i.e.,* growth on supplied C sources) across the 10 locations for each land-use type. This outcome suggests high variation in bacterial growth dynamics across locations (assessed here as differential abundance in response to C addition). However, this finding is generally consistent with the idea that occupancy is a poor predictor of activity and that soil factors, such as initial microbial biomass, may impart ecological controls on activity. Most OTUs that increased in differential abundance in the Enrichment Microcosms also assimilated ^13^C from the same C source in the SIP Microcosms. Many of these were classified in the family *Burkholderiaceae* (Supplemental Data), which also included taxa found to be highly dynamic and to assimilate highly bioavailable C in the DNA-SIP experiment.

While the multi-substrate DNA-SIP experiment we performed provides valuable insight regarding bacterial contributions to soil C-cycling, the experimental design has notable limitations. A foremost limitation is that incubations are performed in microcosms with sieved homogenized soil to minimize experimental variation and improve the sensitivity and specificity of DNA-SIP (Youngblut *et al*., 2018b). Microcosms exclude plant roots, which have major impacts on microbial activity, and so plant effects, such as priming (Kuzyakov *et al*., 2000; Fontaine *et al*., 2003; Pascault *et al*., 2013), are absent from this experiment. In addition, sieving will disrupt soil aggregates and hyphal networks. Fungi were not assessed in this experiment, but fungal activity is a crucial component of soil C-cycling (Koechli *et al*., 2019). *Fungal:Bacterial* ratios are generally higher under low intensity land-use (*e.g.* old-field and forests) than high intensity land-use (*e.g.* cropland) (Bardgett *et al*., 1996; Strickland and Rousk, 2010) and these ratios can affect organic matter turnover rates (Bailey *et al*., 2002; Malik *et al*., 2016). Disruption of hyphae by sieving could also have disproportionate effects on fungi that establish extensive hyphal networks. We expect that fungi interact with bacteria as agents of primary degradation for all C sources tested, and that their metabolic products are available for assimilation by bacteria and other fungi (Koechli *et al*., 2019). Hence, differences in fungal community structure likely contribute to the patterns of C mineralization and bacterial activity we observed. Additional experiments are needed to assess the effects that roots, fungi, and multi-partite interactions have on soil C dynamics. We further expect that C assimilation by many bacteria, particularly at later time points, takes place after microbial processing of added C sources. Our initial goal was not to identify microbes responsible for primary assimilation of specific substrates, but rather to track C from different sources as the bacterial community assimilates it over time. These patterns of assimilation are coupled to mineralization rates both through primary degradation of C sources as well as through secondary transformations of microbial products. Finally, we used DNA yield to indicate relative microbial biomass across land-use because these values tend to correlate strongly (Blagodatskaya *et al*., 2003; Joergensen and Emmerling, 2006; Blagodatskaya and Kuzyakov, 2013; Fornasier *et al*., 2014; Spohn *et al*., 2016). Alternative measurements (Blagodatskaya and Kuzyakov, 2013) such as direct counting (Skinner *et al*., 1952), chloroform fumigation (Vance *et al*., 1987), qPCR (Smith and Osborn, 2009), and PLFA (Frostegård and Bååth, 1996) are widely used as indicators of microbial biomass, though all methods have advantages and disadvantages.

The hypothetical model we propose provides a framework for improving our understanding of microbially mediated terrestrial C cycling. We demonstrated that land-use has profound impacts on the path C takes through the microbial food web, and that these differences are linked to C mineralization dynamics. We also propose that microbial biomass quantity, as assessed by DNA yield, influences C cycling by altering resource channeling between microbes having different life history strategies, with low biomass soils favoring rapid growing organisms that exhibit repeated cycles of boom and bust growth dynamics, while high biomass soils favor microbes adapted for efficient resource acquisition and less dynamic growth cycling.

## Materials and Methods

### Soil collection

The complete set of soils were collected from 10 locations across a 50 km region around Ithaca, NY, USA, with three separate sites within each location corresponding to cropland, old-field, or forest land-use (Table S1; Fig. S1). Soil collection and edaphic property measurements were previously described (Barnett *et al*., 2020) and detailed in SI. All soils were either Inceptisols or Alfisols with similar histories for each land-use. Only soils from the Monkey Run location (Figs. 1, S1A) were used for multi-substrate DNA-SIP. At Monkey Run, wheat was grown on the cropland site, the old-field site was retired from crop production after 1938, and the secondary forest site had been last harvested prior to 1938.

### SIP Microcosms

SIP Microcosms (*i.e.,* soil microcosms for multi-substrate SIP experiments; Fig. 1) consisted of 15 g air dried soil spread evenly across the bottom of 250 ml Erlenmeyer flasks made. Soils were obtained from the Monkey Run location. All five C sources were added to each SIP Microcosm such that 0.4 mg C g^-1^ soil from each source was applied. Xylose, amino acids (mixture), and vanillin were applied as an aqueous solution to achieve 50% water holding capacity. Cellulose and palmitic acid were applied as powder (< 0.25 mm diameter particulates, which approximates the size of particulate organic matter in soil). In each ^13^C-treatment SIP Microcosm, one substrate was >99% ^13^C-labeled and the other four were unlabeled. ^12^C-control SIP Microcosms contained all five substrates unlabeled. SIP Microcosms were destructively sampled based on turnover rates of from Barnett *et al*. (Barnett *et al*., 2021): xylose and amino acids on days 1, 3, 6 and 14; vanillin, cellulose, and palmitic acid on days 6, 14, and 30. Day 0 was sampled before substrates were applied. Rates and cumulative masses of C mineralization were determined from ^12^CO_2_ and ^13^CO_2_ concentrations in the SIP Microcosm headspaces, measured every 1-2 days using GCMS. This design required analysis of 213 SIP Microcosms to accommodate replication and destructive sampling over time. Triplicate SIP Microcosms were made. More detail on SIP Microcosm design is included in SI (Fig. S2).

### Enrichment Microcosms

Enrichment Microcosms (*i.e.,* soil microcosms for the enrichment experiment; Fig. 1) consisted of 3.5 g air dried soil within wells of polystyrene 3x4 well culture plates. Soils were from the three land-use sites from all 10 locations. C sources were added in the same manner as described above, however only one C source was added to each Enrichment Microcosm and no substrates were ^13^C-labeled. Enrichment Microcosms were destructively sampled based on the C source added: xylose, amino acids, and vanillin on days 2 and 4; cellulose and palmitic acid on days 14 and 28. Control Enrichment Microcosms with no added C source were sampled on all four days. This design required analysis of 420 Enrichment Microcosms to accommodate destructive sampling over time (Fig. S3). Replicates for Enrichment Microcosms were operationally impractical.

### DNA extraction

DNA for DNA-SIP was extracted from SIP Microcosm soil using a phenol-chloroform bead-beating method (Griffiths *et al*., 2000). This method produces sufficient DNA quantity and quality for DNA-SIP protocols (Barnett *et al*., 2021). Additional DNA was extracted from SIP Microcosms for bacterial community analyses, hereafter called “unfractionated” DNA. This unfractionated DNA was analyzed directly without isopycnic centrifugation to evaluate whole bacterial community dynamics in SIP Microcosm soils. DNA from Enrichment Microcosm and bulk field soils, as well as the unfractionated DNA, was extracted using the PowerMag Microbiome RNA/DNA Isolation Kit (MO BIO Laboratories Inc., Carlsbad, CA, USA). More details on DNA extractions are found in SI.

### Isopycnic centrifugation of DNA from SIP Microcosms

Isopycnic centrifugation of DNA from the SIP Microcosms was conducted using 4.7 ml tubes and a TLA110 rotor as detailed in SI. A total of 28 fractions (100 μl each) were collected for each CsCl gradient, with buoyant density ranging from 1.67-1.79 g/ml (Pepe-Ranney *et al*., 2016; Barnett *et al*., 2021).

### DNA amplification, sequencing, and read processing

DNA amplification, sequencing, and read processing has been previously described (Barnett *et al*., 2020). In short, the V4 region of the 16S rRNA gene was amplified in triplicate using dual indexed primers based on Kozich *et al*. (Kozich *et al*., 2013). Primer design is detailed in SI. Sequencing was conducted at the Cornell Biotechnology Resource Center (Ithaca, NY, USA) on the Illumina MiSeq platform with the paired end 2 × 250 bp V2 kit. Separately for each amplicon library, paired-end reads were merged using PEAR (Zhang *et al*., 2014), demultiplexed using a custom script, filtered using alignment based quality filtering (Silva SEED database release 132, maximum homopolymer length = 8) with mothur (Schloss *et al*., 2009), and sequences mapping to mitochondria, chloroplasts and Archaea were removed. At this point, all libraries were combined, and operational taxonomic units (OTUs) were clustered at 97% sequence identity using USEARCH (Edgar, 2010). OTU taxonomy was assigned based on SILVA release 132 using the uclust algorithm through QIIME (Caporaso *et al*., 2010). A phylogenetic tree of all OTUs was generated using the PyCogent toolkit (Knight *et al*., 2007). Raw demultiplexed sequences are available at the NCBI Short Read Archive BioProject accession PRJNA686389.

All statistical analyses were conducted using R version 3.6.3. Code, processed sequencing data, and GCMS raw data are available at https://github.com/seb369/FullCyc2.

### Comparing carbon mineralization dynamics across land-use

^13^CO_2_ and ^12^CO_2_ concentrations were measured from SIP Microcosm headspace every 24-48 hours using a Shimadzu GCMS model QP2010S (Shimadzu, Kyoto, Japan), with a Carboxen 1010 PLOT column (Supelco, Bellefonte, PA, USA) as detailed in SI (Barnett *et al*., 2021). We compared the rates of both total C mineralization (^12^C + ^13^C) from the entire substrate mixture and the ^13^C mineralization from each individual ^13^C-labeled substrate across land-use type using linear mixed effects modeling with package nlme (Pinheiro *et al*., 2020). Land-use, days following substrate addition (time), and their interaction were fixed effects and microcosm identity was a random effect. Land-use pairs with significantly differing C mineralization rates at each timepoint were identified from post-hoc pairwise comparisons with package lsmeans (Lenth, 2016) correcting *p-*values for multiple comparisons using Benjamini-Hochberg adjustment. Cumulative C mineralization at the end of the sampling period was compared across land-use types by ANOVA followed by post hoc Tukey tests to identify pairwise differences. All analyses were run separately for each substrate.

### Bacterial community analysis of unfractionated DNA

We rarefied the unfractionated DNA OTU table to an even sampling depth (3302 reads) using function *rarefy_even_depth* from package Phyloseq (McMurdie and Holmes, 2013). Rarefying was used because it is a robust normalization method when samples have uneven sequencing depths and particularly when presence/absence diversity measures are used (Weiss *et al*., 2017). We examined alpha diversity (OTU richness, Shannon’s diversity index, and Pielou’s species evenness) across SIP Microcosms using a linear regression with land-use, time, and their interaction as explanatory variables. Post-hoc pairwise comparisons between land-use were made using package lsmeans (Lenth, 2016), with *p-*values adjusted with the Benjamini-Hochberg procedure. Within each land-use, alpha diversity at each timepoint was compared to the averaged day 0 values using t-tests to determine how substrate addition influenced diversity over time. *P-*values were adjusted with the Benjamini-Hochberg procedure.

We compared bacterial community composition across SIP Microcosms by PERMANOVA using function *adonis* from package vegan (Oksanen *et al*., 2018) with land-use, time, and their interaction as explanatory variables. PERMANOVA were run separately for three beta diversity measures: Bray-Curtis dissimilarity and both weighted and unweighted UniFrac distances. To test the effect of substrate addition on bacterial community composition over time, within each land-use the dissimilarity between each timepoint and day 0 was compared to the dissimilarity between day 0 replicates (*i.e.* bottle effect) via t-tests. *P-*values were adjusted with the Benjamini-Hochberg procedure.

### Identifying ^13^C-incorporators in DNA-SIP experiments

We identified OTUs whose DNA incorporated ^13^C from ^13^C-labeled C sources (*i.e.* incorporators) using multiple-window high-resolution DNA-SIP (MW-HR-SIP) (Youngblut *et al*., 2018b) with the package HTSSIP (Youngblut *et al*., 2018a). We used overlapping buoyant density windows: 1.70-1.73, 1.72-1.75, and 1.74-1.77 g mL^-1^ (Youngblut *et al*., 2018b; Barnett *et al*., 2021). Each window contained between 4-8 fractions with a median of 7. MW-HR-SIP tests for OTUs that are ^13^C-labeled by comparing their differential abundance between corresponding gradient fractions from ^13^C-treatments and identically treated but unlabeled controls (Youngblut *et al*., 2018b). Incorporators were identified as OTUs that are differentially abundant (*p* < 0.05 after adjusting for false discovery and multiple comparisons).

Samples and OTUs were filtered prior to MW-HR-SIP analysis to increase statistical power. Gradient fractions that had less than 1000 reads were removed prior to analysis due to having insufficient reads for statistical analysis (one fraction each from old-field ^13^C-xylose day 3, forest ^13^C-vanillin day 14, and forest ^12^C-control day 30). We also used independent filtering on the basis of sparsity to remove uninformative OTUs by employing a variable sparsity threshold between 0.05-0.50, applied to fractions within the entire 1.70-1.77 g mL^-1^ “heavy” window. For each treatment, the sparsity threshold that provided the greatest information content (the highest number of OTUs with rejected hypotheses) was selected. In addition, OTUs detected in only the ^13^C-treatment or ^12^C-control gradients were removed by filtering prior to analysis because it is impossible to confirm the ^13^C-labeling status of these OTUs.

### Bacterial community analysis in Enrichment Microcosms

We rarefied Enrichment Microcosm OTU tables as before (14503 reads). We again compared bacterial community composition across Enrichment Microcosms by PERMANOVA, with land-use, substrate, timepoint, and their interactions as explanatory variables. PERMANOVA were run separately for Bray-Curtis dissimilarity, weighted, and unweighted UniFrac distances. We were further interested in how the change in bacterial communities due to substrate addition was related to soil parameters. To test this relationship, we first examined change in Pielou’s evenness by subtracting evenness in the no-substrate controls from the evenness in the substrate treatments. Overall loss of evenness was tested with one sided, one sample Wilcoxon tests (µ=0). The relationships between change in evenness and DNA yield were measured using Pearson’s correlation. DNA yield were from the bulk soils to represent initial soil conditions. Second, we determined the change in bacterial community composition due to substrate addition by calculating the weighted UniFrac distance between each substrate-treated Enrichment Microcosm and its corresponding no-substrate control. The relationships between the level of compositional change between treatment and control Enrichment Microcosms and initial DNA yield were then measured using Pearson’s correlation. In all above analyses, *p-*values were adjusted using the Benjamini-Hochberg procedure.

We identified OTUs from each land-use that were enriched by C source additions, relative to buffer only controls, in the Enrichment Microcosms by using DESeq2 (Love *et al*., 2014). DESeq2 analysis used unrarefied OTU tables with sparsity filtering to improve statistical power. For sparsity filtering, OTUs not represented by at least 5 reads in at least 25% of the samples were removed as uninformative prior to statistical analysis. To account for paired soil source locations between treatment and no-substrate controls, the DESeq function used the formula ∼ location + substrate, though OTU enrichment results were returned based on substrate alone. A log 2 fold change threshold of 0.25, one sided Wald test, and adjusted *p-*value cutoff of 0.05 were used to distinguish OTUs enriched by C addition.

## Acknowledgements

We thank Chantal Koechli, Spencer Debenport, Braulio Castillo, and Andrew St. James for their help with experiment implementation and Joseph Yavitt and Ian Hewson for comments on the study and manuscript. Assistance with the linear effects modeling was provided by Lynn Johnson from the Cornell Statistical Consulting Unit.

