## Supplementary material for "Bacterial community dynamics explain carbon mineralization and assimilation in soils of different land-use history": SI

**SUPPLEMENTARY INFORMATION**

**Supplementary Materials and Methods**

*Bulk soil collection*

Soils for bulk soil measurements and Enrichment Microcosms were collected as previously described (Barnett *et al.*, 2020) from 10 locations in a 50 km region around Ithaca, NY, USA (Table S1; Fig. S1). Each location contained a separate cropland, old-field, and forest site. The cropland sites were in active use, cultivating annual field crops following standard management practices for this region. The old-field sites were disused cropland which had not been cultivated for at least 20 years and contained a diverse mix of annual and perennial forbs. The forest sites were secondary growth, dominated by regionally common deciduous trees (*e.g.* maple and beech). Eight of the forest sites are at least 77 years old while the other two are between 30 and 70 years old. Soil types were consistent throughout, with 28 sites classified as Inceptisol and two as Alfisol. All soil samples were collected on the 26th or 27th of October, 2015. In each site, we established a 5 x 5 m sampling plot and collected 20 random soil cores (2.5 cm diameter and 10 cm depth), homogenizing them and sieving to 2 mm. At Monkey Run, the location used for the SIP Microcosms, 80 cores were collected. Aliquots for bulk DNA sequencing were stored at -80˚C while those for soil edaphic property measurements were dried overnight at 105˚C and stored at room temperature. Aliquots for microcosms were stored at 4˚C prior to microcosm construction. Soil edaphic properties were measured from the bulk soil as described previously (Barnett *et al.*, 2020). Soil organic matter content was measured using the loss on ignition method. Soil moisture was measured gravimetrically.

*SIP Microcosms*

Soil microcosms for multi-substrate SIP (SIP Microcosms) were constructed and sampled as previously described (Pepe-Ranney *et al.*, 2016; Barnett *et al.*, 2021). SIP Microcosms consisted of 15 g dry-weight soil from Monkey Run in sterile 250 ml Erlenmeyer flasks capped with rubber stoppers. Five C sources were used in this experiment, representing common plant molecules: xylose (Omicron Biochemicals Inc., South Bend, IN, USA), amino acid mixture (Cambridge Isotope Laboratories Inc., Andover, MA, USA), vanillin (Sigma Isotec, Miamisburg, OH, USA), cellulose (generated in house (Pepe-Ranney *et al.*, 2016)), and palmitic acid (Sigma Isotec). A mixture of all five substrates were added to each SIP Microcosms such that 0.4 mg of carbon from each was applied per gram of dry weight soil. In each ^13^C-treatment SIP Microcosm, one substrate was >99% ^13^C-labeled while the other four had natural background isotope ratios (^12^C). We also made ^12^C-control SIP Microcosms in which all five substrates were unlabeled. Soil sampling days were based on the predicted rates of metabolism of the given substrates as in Barnett *et al.* (Barnett *et al.*, 2021). SIP Microcosms supplied with ^13^C-xylose or ^13^C-amino acids were sampled 1, 3, 6 or 14 days after substrate addition and SIP Microcosms supplied with ^13^C-vanillin, ^13^C-cellulose, or ^13^C-palmitic acid were sampled 6, 14, and 30 days after substrate addition. Soil sampling was destructive, so separate SIP Microcosms were made for each soil sampling timepoint. Soil sampling was accomplished by scooping out all soil from the SIP Microcosms with a sterile spatula and immediately storing soils at -80˚C in Whirl-Pak bags. Day 14 SIP Microcosms for ^13^C-xylose and ^13^C-amino acids and day 30 SIP Microcosms for ^12^C-control, ^13^C-vanillin, ^13^C-cellulose, ^13^C-palmitic acid, and H_2_O control were made in triplicate; all other SIP Microcosms were made in duplicate. Duplicate Day 0 SIP Microcosms were made and sampled before substrate addition. Triplicate no-substrate control SIP Microcosms were made by applying the base salt solution but no substrates. No-substrate control SIP Microcosms were sampled on day 30. We made 213 SIP Microcosms in all.

Prior to substrate addition, microcosms were conditioned at room temperature for 2 weeks. During conditioning, headspace CO_2_ measurements were taken every other day and carbon addition did not occur until after CO_2_ flux from soils had stabilized. To apply the water-soluble substrates (xylose, amino acids, and vanillin), we dissolved appropriate amounts in a 2.9% Murashige Skoog basal salt mixture (Sigma Aldrich M5524). Separate soluble substrate solutions were made up for each land-use so that 0.4 mg of carbon from each substrate was added per gram of dry weight soil while bringing soil moisture up to 50% water holding capacity. Solutions were heated to 65˚C for full dissolution of vanillin then cooled to room temperature and filter sterilized. Solutions were sprayed evenly over the soil within each microcosm using a sterilized mucosal atomization device. Powdered insoluble substrates (cellulose and palmitic acid) were autoclave sterilized. Insoluble substrates were also added by mass to each microcosm such that 0.4 mg of carbon from each was added per gram of dry weight soil. We evenly spread the powdered insoluble substrates over the surface of the microcosm soils, passing through a 250 µm mesh. Microcosms were incubated in the dark at room temperature. Where possible, microcosms were handled in random order to avoid batch effects.

We measured the amount of carbon mineralized from the SIP Microcosm soils by measuring ^12^CO_2_ and ^13^CO_2_ concentration in 250µl of SIP Microcosm headspace. A subset of SIP Microcosms was repeatedly used for headspace sampling throughout the experiment: no-substrate control, ^12^C-control day 30, ^13^C-xylose day 14, ^13^C-amino acid day 14, ^13^C-vanillin day 30, ^13^C-cellulose day 30, and ^13^C-palmitic acid day 30. All three replicates of each were sampled. Headspace samples were collected every day through day 6, then every other day through day 30. ^12^CO_2_ and ^13^CO_2_ concentrations were measured on a Shimadzu GCMS model QP2010S (Shimadzu, Kyoto, Japan), with a Carboxen 1010 PLOT column (Supelco, Bellefonte, PA, USA). After headspace sampling, the headspaces of all SIP Microcosms were flushed with 0.22 µm filtered air. CO_2_ concentration and mass were calculated using standard curves prepared for each sampling timepoint. After soil was removed from SIP Microcosms, we measured the exact volume of each flask with water. SIP Microcosm volume was then used along with the ideal gas law to calculate the mass of ^12^C and ^13^C in the headspace after each 24-48 hour sampling period thereby estimating the rate of carbon mineralization from each substrate (^13^C) and total organic matter pool (^12^C).

*Enrichment Microcosms*

We also ran an enrichment experiment where one C source was added to each microcosm to enrich populations of bacterial taxa growing on these substrates (Enrichment Microcosms). This experiment used soil from each land-use from all 10 regional locations. Enrichment Microcosms were constructed as before, except that they were made in wells of 3x4 well culture plates, consisted of 3.5 g dry weight soil, and sealed with breathable film. Enrichment Microcosm assignments were randomized across all the culture plates to avoid batch effects. For these enrichments, only one substrate was added to each Enrichment Microcosm and no substrates were isotopically labeled. Substrates were prepared and applied to the Enrichment Microcosms as before with a final carbon added mass of 0.4 mg per gram of dry weight soil. Prior to substrate addition, Enrichment Microcosms were conditioned at room temperature for 2 weeks. Sterile base salt solution was sprayed over Enrichment Microcosms containing insoluble substrates, while soluble substrates were applied dissolved within this solution. In all cases, enough base salt solution was applied to bring soil moisture to 50% water holding capacity. For each timepoint and soil, a control Enrichment Microcosm was created with no substrate added, but similarly sprayed with the base salt solution. Soils treated with xylose, amino acids, and vanillin were sampled on days 2 and 4 while those treated with cellulose and palmitic acid were sampled on days 14 and 28. No-substrate controls were sampled at the same timepoints. Soil sampling was conducted as before. No replicate Enrichment Microcosms were used in this case. We made 420 enrichment Enrichment Microcosms in total.

*DNA extraction*

DNA destined for isopycnic centrifugation was extracted from 0.25 g of SIP Microcosm soil using a modified Griffiths phenol-chloroform method (Griffiths *et al.*, 2000). Five replicate DNA extractions were performed for old-field and forest soils to recover over 5 µg of total DNA. Due to lower biomass, ten replicate extractions were performed for all cropland soil samples to recover 5 µg of total DNA. Replicate DNA extracts were pooled. DNA fragments less than 4 kb were removed using BluePippin (Sage Science, Beverly, MA) using standard protocol. Size selection was done to avoid gradient smearing from small DNA fragments that would not reach equilibrium during the centrifugation time we used (Youngblut and Buckley, 2014). DNA concentration was quantified using the Quant-iT PicoGreen assay kit (cat# P7589, Thermo Fisher Scientific, Waltham MA, USA). If the resulting product contained less than 5 µg of DNA, extraction and size selection was repeated as above and then pooled with the initial extract.

DNA was extracted from bulk soils, Enrichment Microcosms, and SIP Microcosms not destined for isopycnic centrifugation (unfractionated DNA) as previously described (Barnett *et al.*, 2020) . This DNA was extracted with the PowerMag Microbiome RNA/DNA Isolation Kit (cat# 27 500–4-EP, MO BIO Laboratories Inc. Carlsbad CA, USA) according to manufacturer's specifications except that homogenization was performed by bead beating for 1 min using a Mini-Beadbeater-96 (cat# 1001, Biospec Products, Bartlesville OK, USA).

*Isopycnic centrifugation of DNA from SIP Microcosms*

Isopycnic centrifugation of DNA from the SIP Microcosms was conducted as previously described (Pepe-Ranney *et al.*, 2016; Barnett *et al.*, 2021). CsCl gradients consisted of 4.7 ml OptiSeal ultracentrifuge tubes (Cat # 361621, Beckman Coulter, Brea CA) containing 7.5766 g (4.3 ml) of CsCl gradient solution (1.762 g/ml CsCl, 15 mM Tris-HCl (pH 8.0), 15 mM KCl, and 15 mM EDTA). To this we added 5 µg of aqueous DNA followed by enough 1X TE buffer filler to bring the added fluid to 450 µl. Up to 8 gradients were centrifuged per run at 55,000 rpm and 20˚C for 66 hours on a Beckman Coulter Optima MAX-E ultracentrifuge (Beckman Coulter, Brea CA, USA) with a TLA-110 fixed angle rotor.

Immediately after centrifugation, gradients were fractionated *via* water displacement by pumping distilled water into the top of the tube using a syringe pump in 100 µl increments. Displaced fractions dripped out the bottom of the ultracentrifuge tube, which was punctured with a sterile 18G needle. The refractive index was measured for each fraction using a Reichert AR200 digital refractometer (Reichert Technologies, Buffalo NY, USA) immediately after it was collected to avoid error due to evaporation. From the refractive index we calculated the buoyant density as previously described (Buckley *et al.*, 2007). About 28 fraction were collected for each gradient, with buoyant density ranges from about 1.67-1.79 g/ml. Fractions were desalted using AMPure XP magnetic beads (Cat #A63882, Beckman Coulter, Brea, CA, USA).

*DNA amplification and sequencing*

The V4 region of the 16S rRNA gene was amplified using dual indexing primers as previously described (Barnett *et al.*, 2020). Two primer sets were used, which were both versions of the 515f-806r primers developed by Kozich et al. (Kozich *et al.*, 2013). For bulk soil, unfractionated DNA, and DNA-SIP fractions, the forward 515f primer was as reported in Kozich et al., while the reverse 806r primer, had the pad 5′-AATGTTTTAA-3′, linker 5′-TG-3′, and 16S rRNA specific region 5′-GTGYCAGCMGCMGCGGTRA-3′ (Barnett *et al.*, 2020), with indexing from Kozich et al. For the Enrichment Microcosm soils, the unmodified Kozich et al. primers were used. Primers were changed for the Enrichment Microcosms to improve sequence quality.

Polymerase chain reaction (PCR) amplification, cleanup, and sequencing was performed as previously described (Barnett *et al.*, 2020), though with different DNA template concentration based on DNA source. For DNA from field soil, Enrichment Microcosms, and unfractionated SIP Microcosms, 2 µl of template diluted 1:10 was used per reaction. For DNA-SIP fractions, template was added in volumes or dilutions such that 1 ng of DNA was used per reaction; however, if the volume required to get 1 ng exceeded 5 µl, only 5 µl of template was used. Amplification was performed in 25 µl triplicate reactions with 13.1 µl of Q5 hot start high-fidelity master mix (cat# M0494L, New England Biolabs, Ipswich, MA, USA) mixed 1:0.025 v/v with 200 4X Quant-iT PicoGreen reagent (cat# P7589, Thermo Fisher Scientific, Waltham, MA, USA), 2.5 µl mixed 10X primers, and enough PCR grade water for 25 µl total volume. PCR conditions were 95˚C for 2 min followed by 30 cycles of 95˚C for 20 sec, 55˚C for 15 sec and 72˚C for 10 sec, and followed 72˚C for 5 min. Triplicate PCR products were pooled and then normalized with the Invitrogen SequalPrep Normalization Plate Kit (cat# A1051001, Thermo Fisher Scientific, Waltham, MA, USA). Up to 192 samples were pooled together per amplicon library and sequenced at the Cornell Biotechnology Resource Center (Ithaca, NY, USA) on the Illumina MiSeq platform with the paired end 2 × 250 bp V2 kit.

*Sequence processing*

Sequence processing was conducted as previously described (Barnett *et al.*, 2020). Separately for each amplicon library, paired-end reads were merged using PEAR (Zhang *et al.*, 2014), demultiplexed using a custom script, filtered using alignment-based quality filtering (Silva SEED database release 132, maximum homopolymer length = 8) with Mothur (Schloss *et al.*, 2009), and sequences mapping to mitochondria, chloroplasts and Archaea were removed. At this point, all libraries were combined. We clustered operational taxonomic units (OTUs) at 97% sequence identity and removed chimeras using USEARCH (Edgar, 2010). OTU taxonomy was assigned based on SILVA release 132 using the uclust algorithm through QIIME (Caporaso *et al.*, 2010). A phylogenetic tree of all OTUs was generated using the PyCogent toolkit (Knight *et al.*, 2007). Raw demultiplexed sequences are available at the NCBI Short Read Archive.

**Supplementary Results**

*Carbon mineralization dynamics*

To assess carbon (C) mineralization dynamics, we measured C mineralization from the total organic matter (*i.e.* total C = ^12^C + ^13^C) and from the individual substrates (^13^C) in the headspace of replicate SIP Microcosms. Results from individual substrates are found in the main text. Total C mineralization rate and cumulative total C mineralized differed with respect to land-use, time, and their interaction (Rate: Table S2, linear mixed effects model, *p*-value < 0.05; Cumulative: Table S3, ANOVA, *p*-value < 0.05). Total C mineralization in cropland soil differed from both old-field and forest soils throughout most of the experiment, while old-field and forest C mineralization differed only at their peaks on day 2 (Fig. S2). Cumulative total C mineralized varied across all three land-use regimes (forest > old-field > cropland; Table S3, Fig. S3).

**Table S1:** Soil metadata from the 10 locations. All 10 locations were used for Enrichment Microcosms and Monkey Run was used for SIP Microcosms. SOM is the percent soil organic matter measured with loss on ignition. Moisture is the percent soil moisture measured gravimetrically. DNA yield is the concentration of DNA recovered from 0.25g of soil

| **Location** | **Land-use** | **Date**  **collected** | **Latitude**  (˚N) | **Longitude**  (˚W) | **Current**  **crop** | **Soil**  **class** | **SOM (%)** | **Moisture**  **(%)** | **DNA yield**  **(ng/µl)** |
| --- | --- | --- | --- | --- | --- | --- | --- | --- | --- |
| Bald Hill | Cropland | 10/26/15 | 42.35870 | 76.34999 | Soybeans | Inceptisol | 11.75 | 28.5 | 39.765 |
| Bald Hill | Forest | 10/26/15 | 42.35822 | 76.37527 |  | Inceptisol | 13.19 | 30.1 | 64.906 |
| Bald Hill | Old-field | 10/26/15 | 42.35358 | 76.37195 |  | Inceptisol | 9.97 | 38.5 | 41.075 |
| Caldwell Fields | Cropland | 10/27/15 | 42.44894 | 76.45752 | Buckwheat | Inceptisol | 3.08 | 15.2 | 4.766 |
| Caldwell Fields | Forest | 10/27/15 | 42.45148 | 76.46150 |  | Inceptisol | 12.47 | 30.9 | 59.901 |
| Caldwell Fields | Old-field | 10/27/15 | 42.45062 | 76.45900 |  | Inceptisol | 6.60 | 26.7 | 61.298 |
| Carter Creek | Cropland | 10/27/15 | 42.31995 | 76.66270 | Barley | Inceptisol | 5.97 | 20.9 | 18.764 |
| Carter Creek | Forest | 10/27/15 | 42.33502 | 76.66756 |  | Inceptisol | 15.73 | 29.1 | 53.043 |
| Carter Creek | Old-field | 10/27/15 | 42.32268 | 76.66626 |  | Inceptisol | 11.25 | 30.7 | 76.509 |
| Edwards Lake Cliff Preserve | Cropland | 10/26/15 | 42.51730 | 76.49904 | Hay | Alfisol | 4.93 | 20.6 | 19.965 |
| Edwards Lake Cliff Preserve | Forest | 10/26/15 | 42.52263 | 76.51993 |  | Inceptisol | 11.97 | 30.9 | 54.607 |
| Edwards Lake Cliff Preserve | Old-field | 10/26/15 | 42.52375 | 76.51796 |  | Inceptisol | 10.37 | 34.6 | 41.645 |
| McGowan Woods | Cropland | 10/27/15 | 42.44884 | 76.45116 | Wheat | Inceptisol | 3.29 | 18.3 | 9.834 |
| McGowan Woods | Forest | 10/27/15 | 42.44713 | 76.45036 |  | Inceptisol | 7.92 | 29.1 | 68.800 |
| McGowan Woods | Old-field | 10/27/15 | 42.44835 | 76.45488 |  | Inceptisol | 4.68 | 20.4 | 17.866 |
| Monkey Run | Cropland | 10/27/15 | 42.46888 | 76.43430 | Wheat | Inceptisol | 4.85 | 19.0 | 8.294 |
| Monkey Run | Forest | 10/27/15 | 42.47114 | 76.42986 |  | Inceptisol | 11.31 | 23.2 | 60.650 |
| Monkey Run | Old-field | 10/27/15 | 42.46949 | 76.43089 |  | Inceptisol | 8.17 | 28.2 | 60.644 |
| Mount Pleasant | Cropland | 10/26/15 | 42.45980 | 76.38463 | Rye clover | Inceptisol | 6.05 | 22.9 | 32.446 |
| Mount Pleasant | Forest | 10/26/15 | 42.46718 | 76.38340 |  | Inceptisol | 9.81 | 30.0 | 72.924 |
| Mount Pleasant | Old-field | 10/26/15 | 42.46226 | 76.37859 |  | Inceptisol | 12.75 | 32.7 | 102.073 |
| Musgrave Farm | Cropland | 10/26/15 | 42.72776 | 76.65459 | Corn | Inceptisol | 4.89 | 18.7 | 4.432 |
| Musgrave Farm | Forest | 10/26/15 | 42.72883 | 76.64992 |  | Inceptisol | 12.98 | 28.3 | 54.671 |
| Musgrave Farm | Old-field | 10/26/15 | 42.73157 | 76.66258 |  | Alfisol | 10.66 | 28.6 | 60.027 |
| Polson Preserve | Cropland | 10/26/15 | 42.43158 | 76.38955 | Rye clover | Inceptisol | 5.43 | 19.9 | 21.027 |
| Polson Preserve | Forest | 10/26/15 | 42.42597 | 76.39698 |  | Inceptisol | 8.65 | 29.3 | 74.413 |
| Polson Preserve | Old-field | 10/26/15 | 42.42818 | 76.40222 |  | Inceptisol | 11.03 | 33.3 | 65.229 |
| Slaterville 600 | Cropland | 10/26/15 | 42.41070 | 76.33729 | Hay | Inceptisol | 10.93 | 30.7 | 72.756 |
| Slaterville 600 | Forest | 10/26/15 | 42.41757 | 76.32956 |  | Inceptisol | 19.52 | 36.3 | 82.298 |
| Slaterville 600 | Old-field | 10/26/15 | 42.41906 | 76.33344 |  | Inceptisol | 10.00 | 32.1 | 59.115 |

**Table S2:** Linear mixed effects models evaluating the effect of land-use, time, and their interaction on rates of CO_2_ production. Total carbon indicates overall rates of CO_2_-C production (^12^C + ^13^C), while mineralization rates for individual substrates are indicated from ^13^CO_2_.

| **Substrate** | **Factor** | **numDF** | **denDF** | ***F*-value** | ***p*-value** |
| --- | --- | --- | --- | --- | --- |
| Total Carbon | Intercept | 1 | 774 | 38978.14 | < 0.0001 |
|  | land-use | 2 | 51 | 237.08 | < 0.0001 |
|  | day | 18 | 774 | 1222.09 | < 0.0001 |
|  | land-use:day | 36 | 774 | 368.63 | < 0.0001 |
| Cellulose | Intercept | 1 | 108 | 2895.3768 | < 0.0001 |
|  | land-use | 2 | 6 | 8.809208 | 0.0164 |
|  | day | 18 | 108 | 40.509035 | < 0.0001 |
|  | land-use:day | 36 | 108 | 11.308458 | < 0.0001 |
| Xylose | Intercept | 1 | 60 | 3611.76252 | < 0.0001 |
|  | land-use | 2 | 6 | 1.973871 | 0.219 |
|  | day | 10 | 60 | 560.506825 | < 0.0001 |
|  | land-use:day | 20 | 60 | 199.296594 | < 0.0001 |
| Amino acids | Intercept | 1 | 60 | 1708.45467 | < 0.0001 |
|  | land-use | 2 | 6 | 5.29034 | 0.0474 |
|  | day | 10 | 60 | 288.64455 | < 0.0001 |
|  | land-use:day | 20 | 60 | 82.248169 | < 0.0001 |
| Vanillin | Intercept | 1 | 108 | 3919.29109 | < 0.0001 |
|  | land-use | 2 | 6 | 160.673143 | < 0.0001 |
|  | day | 18 | 108 | 320.220176 | <0.0001 |
|  | land-use:day | 36 | 108 | 118.151045 | <0.0001 |
| Palmitic acid | Intercept | 1 | 108 | 962.869682 | <0.0001 |
|  | land-use | 2 | 6 | 1.440048 | 0.308 |
|  | day | 18 | 108 | 33.269555 | <0.0001 |
|  | land-use:day | 36 | 108 | 2.068988 | 0.00218 |

**Table S3:** ANOVA of cumulative mineralization of total carbon (^12^C + ^13^C) and carbon from individual substrates (^13^C) by the end of the experimental periods (Day).

| **Substrate** | **Day** | ***df*** | ***F*-value** | ***p*-value** |
| --- | --- | --- | --- | --- |
| Total carbon | 14 | 2 | 93.068 | < 0.001 |
| Total carbon | 30 | 2 | 367.1709 | < 0.001 |
| Xylose | 14 | 2 | 2.3327 | 0.178 |
| Amino acids | 14 | 2 | 7.486 | 0.0234 |
| Vanillin | 30 | 2 | 201.952 | < 0.001 |
| Cellulose | 30 | 2 | 7.376 | 0.0242 |
| Palmitic acid | 30 | 2 | 1.7389 | 0.2537 |

**Table S4:** Linear regression analysis of alpha diversity metrics (OTU richness, Shannon’s index, and Pielou’s evenness) across land-use type, time since substrate addition (day), and their interaction across the SIP Microcosms (unfractionated DNA).

| **Metric** | **Factor** | ***df*** | ***F-*value** | ***p*-value** |
| --- | --- | --- | --- | --- |
| Richness | land-use | 2 | 646.375 | < 0.001 |
|  | day | 5 | 25.24 | < 0.001 |
|  | land-use:day | 10 | 16.365 | < 0.001 |
| Shannon | land-use | 2 | 676.537 | < 0.001 |
|  | day | 5 | 76.753 | < 0.001 |
|  | land-use:day | 10 | 55.694 | < 0.001 |
| Evenness | land-use | 2 | 488.877 | < 0.001 |
|  | day | 5 | 73.231 | < 0.001 |
|  | land-use:day | 10 | 55.133 | < 0.001 |

**Table S5:** Changes in community composition over time vary with initial soil DNA yield in Enrichment Microcosms. Change in community composition was assessed relative to water-only controls as measured by change in Pielou’s evenness and weighted UniFrac distance. Two tests were run: one sided - one sample Wilcoxon test to compare loss of evenness (*µ* = 0) and Pearson’s correlation with respect to initial DNA yield.

|  |  |  | **Wilcoxon test** | | **Pearson’s correlation to DNA yield** | |
| --- | --- | --- | --- | --- | --- | --- |
| **Metric** | **Day** | **Substrate** | ***V*** | **Adjusted**  ***p-*value** | ***r*** | **Adjusted**  ***p-*value** |
| Change in evenness | 2 | Xylose | 177 | 0.2098 | 0.4805 | 0.036 |
|  | 2 | Amino acids | 70 | < 0.001 | 0.6555 | < 0.001 |
|  | 2 | Vanillin | 15 | < 0.001 | 0.5498 | 0.0082 |
|  | 14 | Cellulose | 210 | 0.3277 | 0.3028 | 0.5194 |
|  | 14 | Palmitic acid | 172 | 0.2098 | 0.1253 | 1 |
|  | 4 | Xylose | 215 | 0.6086 | 0.0266 | 1 |
|  | 4 | Amino acids | 98 | 0.0058 | 0.3834 | 0.1825 |
|  | 4 | Vanillin | 18 | < 0.001 | 0.5857 | 0.0034 |
|  | 28 | Cellulose | 230 | 0.7527 | -0.1949 | 1 |
|  | 28 | Palmitic acid | 249 | 0.7527 | 0.0054 | 1 |
| Weighted UniFrac | 2 | Xylose |  |  | -0.3919 | 0.161 |
|  | 2 | Amino acids |  |  | -0.5862 | 0.0033 |
|  | 2 | Vanillin |  |  | -0.5794 | 0.004 |
|  | 14 | Cellulose |  |  | -0.1308 | 1 |
|  | 14 | Palmitic acid |  |  | -0.3894 | 0.1839 |
|  | 4 | Xylose |  |  | -0.5763 | 0.0043 |
|  | 4 | Amino acids |  |  | -0.2342 | 1 |
|  | 4 | Vanillin |  |  | -0.5619 | 0.0062 |
|  | 28 | Cellulose |  |  | -0.4051 | 0.1463 |
|  | 28 | Palmitic acid |  |  | -0.2616 | 0.8526 |

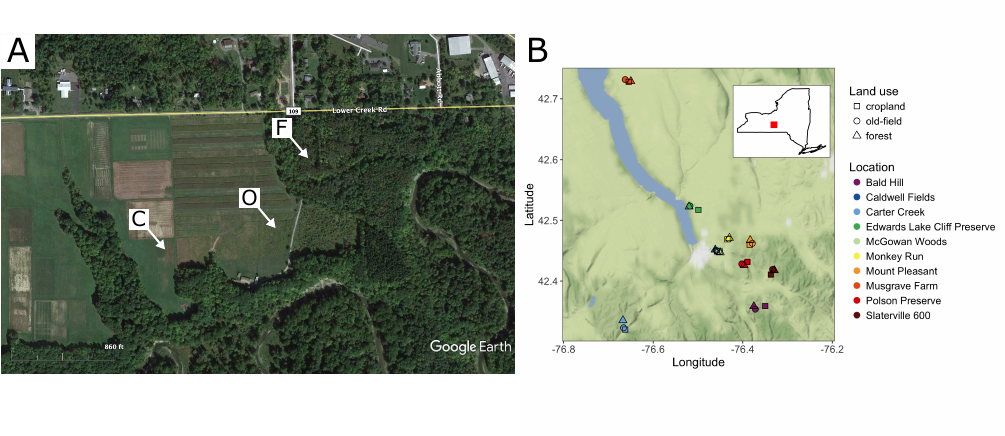

**Figure S1:** Locations of collection sites for soil microcosms. A) Satellite view of the Monkey Run sites used for the SIP Microcosms. Land-use regimes indicated by letters: C = cropland, O = old-field, and F = forest. Image taken in June 2016. B) Locations of all 30 sites used for the Enrichment Microcosms. Land-use indicated by point shape and location indicated by color. Inset is the location of this 50 x 50 km region in New York State. Map tile from Stamen Maps (maps.stamen.com). Both maps oriented with North at the top.

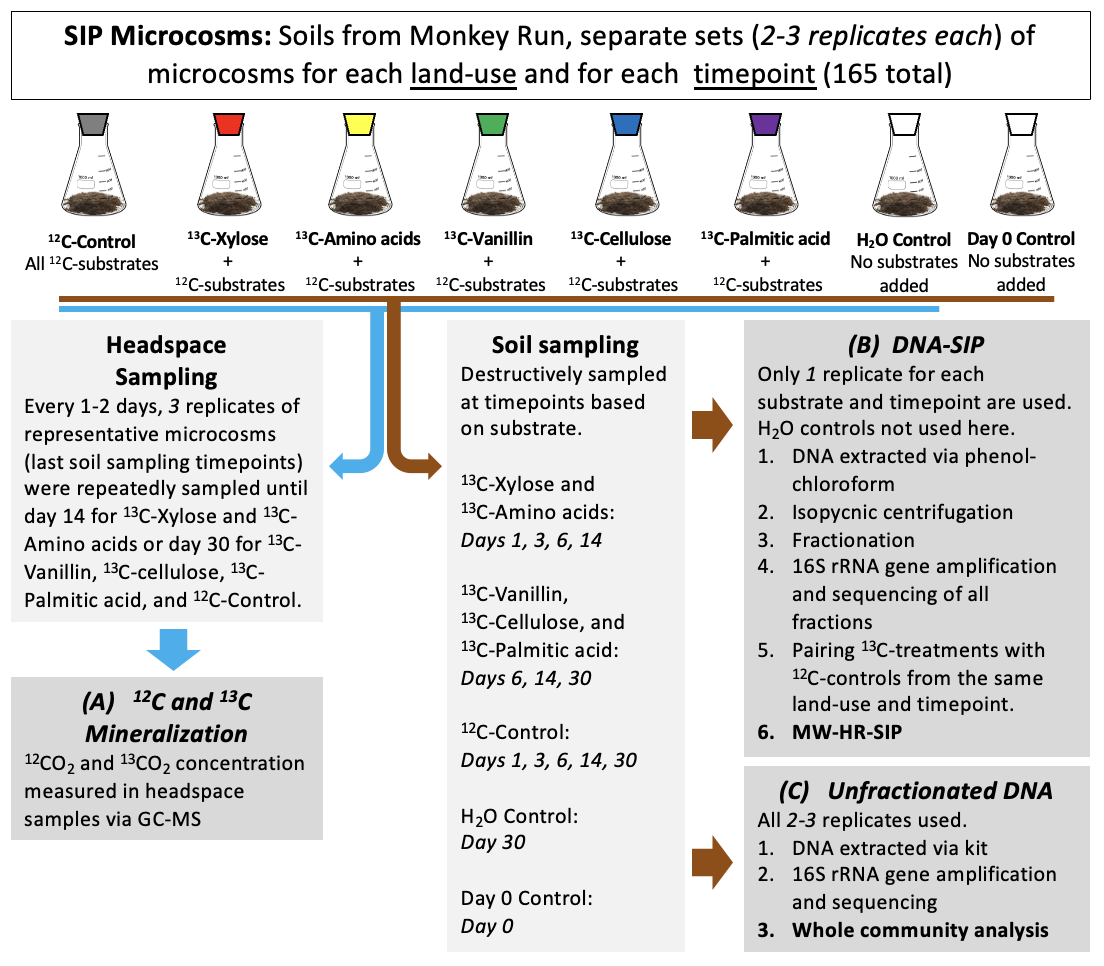

**Figure S2:** Diagram of SIP Microcosm design, sampling, and measurements. These microcosms consisted of 15 g dry weight soil in 250 ml Erlenmeyer flasks sealed with a rubber stopper. SIP Microcosms were made with soils from Monkey Run and separately for each land-use. Separate sets of replicate microcosms were then made for each ^13^C-labeled substrate and substrate-specific timepoints. For ^12^C-Controls and ^13^C-treatments, all 5 substrates were added to each microcosm in equivalent amounts of carbon. Day 14 microcosms for ^13^C-Xylose and ^13^C-Amino acids and Day 30 microcosms for ^12^C-Control, ^13^C-Vanillin, ^13^C-Cellulose, ^13^C-Palmitic acid, and H_2_O control were made in triplicate; all other microcosms were made in duplicate. Output data includes *(A)* quantity of ^12^C and ^13^C mineralized from representative microcosms over time; *(B)* list of OTUs that have evidence of ^13^C incorporation into DNA from each substrate, at each timepoint, and from each land-use; and *(C)* table with abundances of all OTUs in each replicate microcosm from each timepoint, from each treatment, from each land-use.

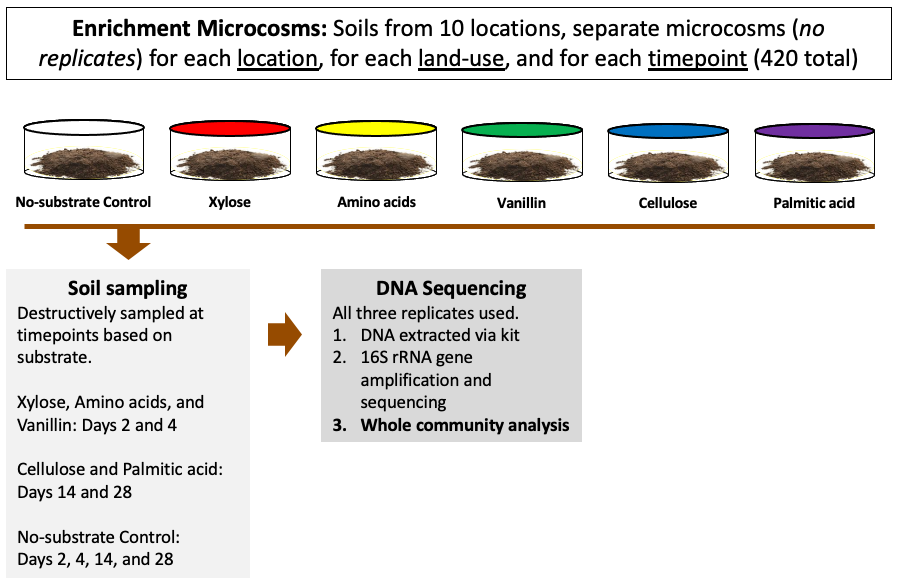

**Figure S3:** Diagram of Enrichment Microcosm design, sampling, and measurements. These microcosms consisted of 3.5 g dry weight soil in 6-well culture plates topped with breathable film. Enrichment Microcosms were made with air dried (two weeks) soils representing each land-use (*n* = 3) from each location (*n* = 10) for a total of 30 sites. For each of the 30 sites, separate microcosms were made for each substrate (*n* = 6, five C sources plus no-carbon control) and each time point (*n* = 2) to accommodate destructive sampling. Replication was made at the level of land-use across the region (10 sites per land-use type), but replication was not performed within each site. This design produced a total of 420 samples. Output data was a table with abundances of all OTUs in each microcosm from each timepoint, from each treatment, from each land-use, from each location.

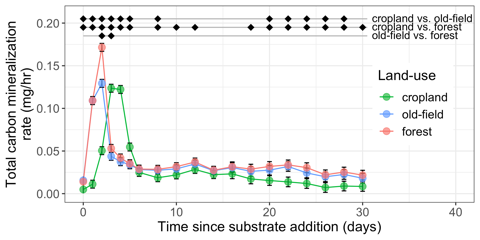

**Figure S4:** Total carbon (^12^C + ^13^C) mineralization rate varied across all three land-use regimes and time (Table S2, linear mixed effect model, *p-*value < 0.05). Statistically significant post-hoc pairwise comparisons (*p-*value < 0.05) are indicated by diamonds at the top with pairwise comparisons between land-use indicated by the horizontal line. Error bars indicate standard error.

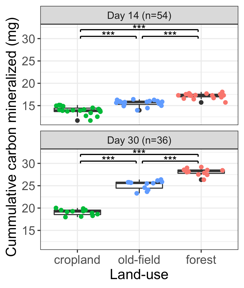

**Figure S5:** Cumulative total carbon (^12^C + ^13^C) mineralized varied across land-use regimes based on measurements for both day 14 and 30 (ANOVA, *p-*value < 0.05). Significant pairwise comparisons indicated by brackets (Tukey test, *** *p-*value < 0.001). *n* is the number of samples used in the analysis.

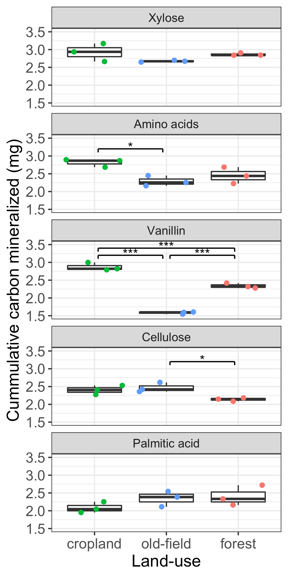

**Figure S6:** Cumulative carbon mineralized from each substrate (^13^C) across land-use regimes at the end of the experimental periods (day 14 for xylose and amino acids, day 30 for vanillin, cellulose, and palmitic acid). Significant pairwise comparisons indicated by brackets (Tukey tests, * *p-*value < 0.05, *** *p-*value < 0.001).

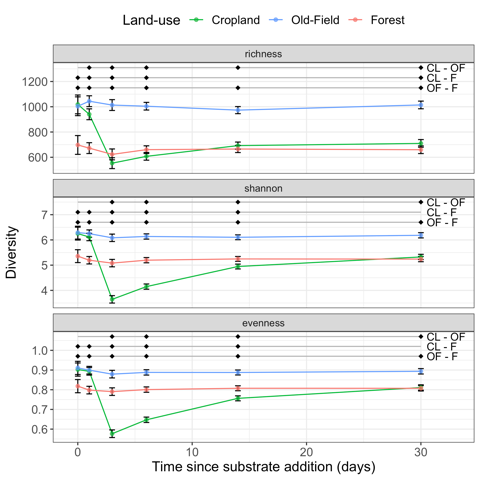

**Figure S7:** Bacterial community alpha diversity (OTU richness, Shannon’s index, and Pielou’s evenness) in SIP Microcosms (unfractionated DNA) varied across land-use regime over time following substrate addition. Statistically significant pairwise differences (*p-*values < 0.05) between land-use regimes indicated by black diamonds at the top of plots with land-use pairs indicated by the horizontal lines (CL = cropland, OF = old-field, F = forest). Error bars indicate standard error.

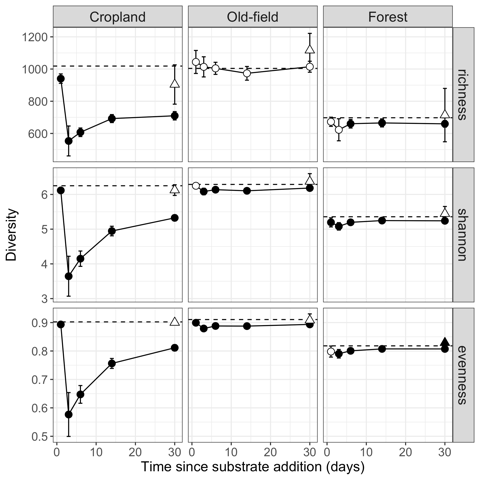

**Figure S8:** Bacterial community alpha diversity (OTU richness, Shannon’s index, and Pielou’s evenness) in SIP Microcosms (unfractionated DNA) expressed relative to initial diversity of soils prior to C addition (indicated by dashed line). Solid symbols indicate time points that differ significantly from initial levels of diversity (t-test, p-value < 0.05), while non-significant points are indicated with open symbols (*p-*value > 0.05). Triangles represent the no-substrate control microcosms (buffer only) sampled on day 30. Error bars indicate standard error.

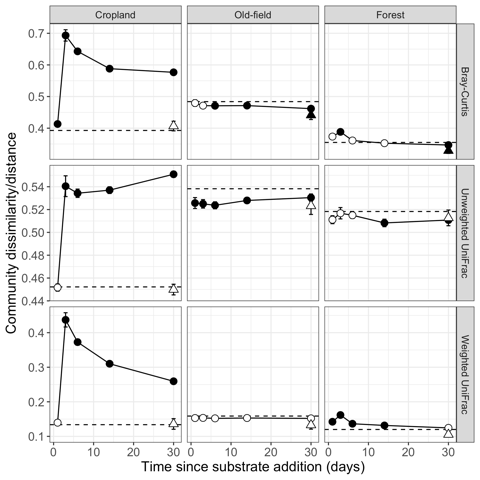

**Figure S9:** Differences in bacterial beta diversity in SIP Microcosms (unfractionated DNA) for each time point following carbon addition, expressed relative to initial community composition. Beta diversity was measured with Bray-Curtis dissimilarity, unweighted UniFrac distance, and weighted UniFrac distance. Initial beta diversity (dashed line) is calculated from comparisons between replicates prior to C addition. Time points that differ significantly from the initial community are indicated by solid symbols (t-test, *p-*value < 0.05) and non-significant comparisons are indicated by open symbols (*p-*value > 0.05). Triangle symbols represent the no-substrate control (buffer only) sampled on day 30. Error bars indicate standard error.

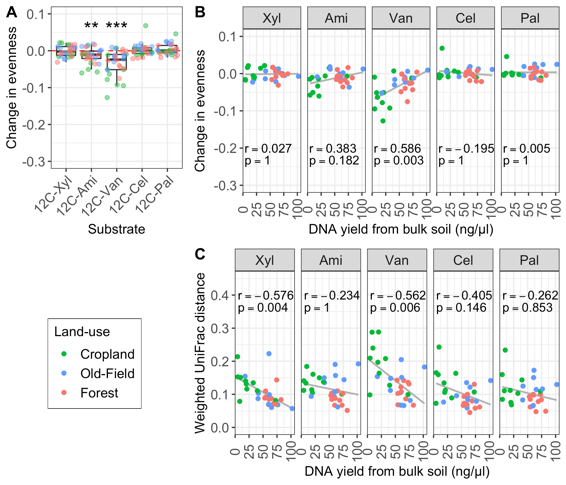

**Figure S10:** Differences in bacterial diversity between substrate added and water-only control Enrichment Microcosms at late timepoints (*i.e.* xylose, amino acid, vanillin: day 4; cellulose and palmitic acid: day 28) were weakly associated with initial microbial population size across 30 sites. DNA yields (ng µl^-1^) from the bulk soils are a proxy for initial microbial population. A) Overall, bacterial communities in soils where amino acids or vanillin were added had significantly less evenness than soils without substrate added (Pielou’s evenness; one-sample, one-sided Wilcoxon test, *** adjusted *p-*value < 0.001). B) Correlations between DNA yield and the change in evenness between substrate added and water-only control microcosms. A positive correlation indicates that smaller initial population led to a loss in evenness over time. C) Correlations between DNA yield and betadiversity (weighted UniFrac distance) between substrate added and water-only control microcosms. A negative correlation indicates that smaller initial population exhibited greater change in betadiversity over time relative to water-only controls. For both (B) and (C) Pearson’s r and Benjamini-Hochberg adjusted *p-*value displayed in graphs and the line represents the linear relationships. Substrates are abbreviated: Xyl = xylose, Ami = amino acids, Van = vanillin, Cel = cellulose, Pal = palmitic acid.

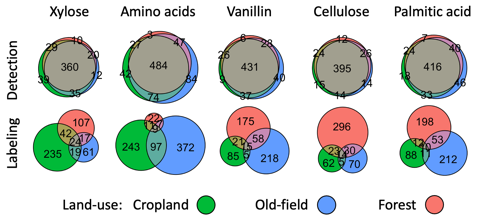

**Figure S11:** Most incorporators were detected (Detection) in all three land-use types but they were ^13^C-labeled (Labeling) in only one land-use type. Detection was defined as passing sparsity filtering prior to MW-HR-SIP.

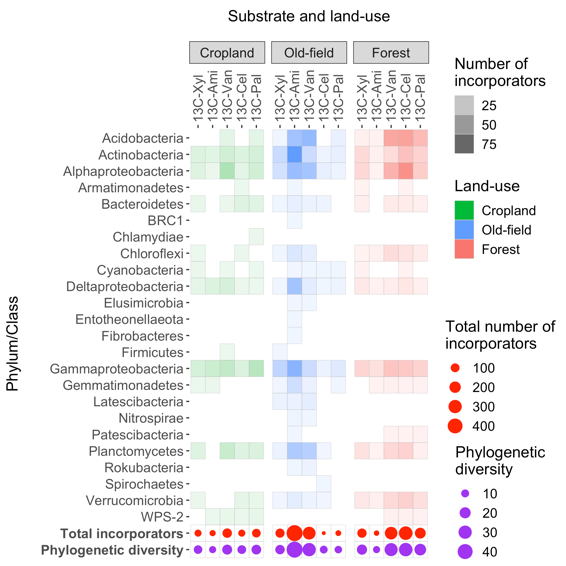

**Figure S12:** Taxonomic makeup of the OTUs incorporating carbon from each substrate at late timepoints (*i.e.* labeled after the day of peak mineralization: days 6 and 14 for xylose and amino acids, days 14 and 30 for vanillin, and day 30 for cellulose and palmitic acid). Taxonomy is reported at the phylum level, with class level reported for *Proteobacteria*. Note that the order *Betaproteobacteriales* is classified within the class *Gammaproteobacteria*. Total number and Faith’s phylogenetic diversity of incorporators of carbon from each substrate in each land-use is indicated by the size of the circles at the bottom. Substrates are abbreviated: 13C-Xyl = xylose, 13C-Ami = amino acids, 13C-Van = vanillin, 13C-Cel = cellulose, 13C-Pal = palmitic acid.

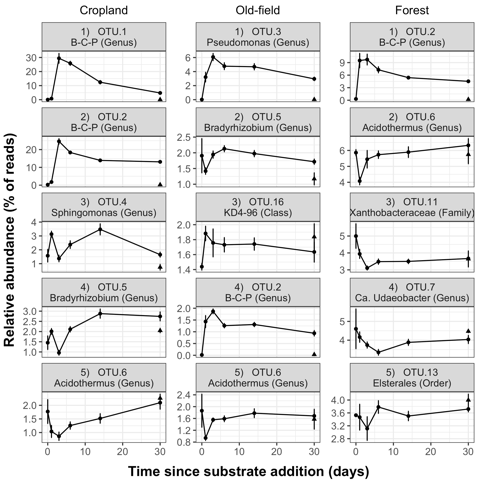

**Figure S13:** Trends in relative abundances for the top 5 most abundant OTUs from all sampling days in each of the three land-use regimes. The OTU classification is given below the rank and OTU name. *B-C-P* represents the *Burkholderia-Caballeronia-Paraburkholderia* genus used by SILVA version 132. Relative abundance is calculated as the percentage of rarefied reads assigned to that OTU. Error bars indicate standard error.
